# A unified neural account of contextual and individual differences in altruism

**DOI:** 10.1101/2022.05.29.493933

**Authors:** Jie Hu, Arkady Konovalov, Christian C. Ruff

**Author notes:** Corresponding authors: Jie Hu and Christian C. Ruff, and. **Author Contributions:** J.H., A.K., and C.C.R. designed research; J.H. and A.K. performed research; J.H., A.K., and C.C.R. analyzed data; and J.H., A.K., and C.C.R. wrote the paper.

## Abstract

Altruism is critical for cooperation and productivity in human societies, but is known to vary strongly across contexts and individuals. The origin of these differences is largely unknown, but may in principle reflect variations in different types of neurocognitive processes that temporally unfold during altruistic decision making (ranging from initial perceptual processing via value computations to final integrative choice mechanisms). Here, we address this question by examining altruistic choices in different inequality contexts with computational modeling and EEG. Our results show that across all contexts and individuals, wealth distribution choices recruit a similar late decision process evident in model-predicted evidence accumulation signals over parietal regions. Contextual and individual differences in behavior related instead to initial processing of stimulus-locked inequality-related value information in centroparietal and centrofrontal sensors, as well as to gamma-band synchronization of these value-related signals with parietal response-locked evidence-accumulation signals. Our findings suggest separable biological bases for individual and contextual differences in altruism and emphasize that these reflect differences in processing of choice-relevant information.

## Introduction

Altruism – incurring own costs to benefit others - is fundamental for cooperation and productivity in human societies (de Waal, 2008; Piliavin and Charng, 1990). It not only plays crucial roles in shaping social political ideology and welfare (e.g., via tax policies and charity) (Bechtel et al., 2018; Offer and Pinker, 2017) but is also essential for collective management of challenging situations, such as political, financial, and public health crises. While altruism is thought to be a stable behavioral tendency shaped by the evolutionary advantages of the ability to cooperate, it is unclear why this tendency varies so strongly across individuals, contexts, and cultures (Bester and Güth, 1998; W D Hamilton, 1964; W.D. Hamilton, 1964; Lebow, 2018; Piliavin and Charng, 1990). Is this because altruism is governed by unitary psychological mechanisms that are engaged in a different manner in different situations or different people (Tricomi et al., 2010)? Or are there fundamentally different types of altruistic actions, guided by different psychological processes, that are triggered differentially in different contexts (Hein et al., 2016)?

From a neurobiological perspective, both these possibilities appear plausible. On the one hand, all altruistic actions necessitate the ability to override self-interest, a parsimonious brain mechanism (Bester and Güth, 1998) that is thought to be facilitated more or less by different contexts and that could be expressed to different degrees in different people (Morishima et al., 2012; Trivers, 1971). On the other hand, empirical observations suggest that altruism varies with factors such as others’ previous actions (e.g. empathy-based vs. reciprocity-based altruism) or their perceived similarity (e.g. social distance) (Hein et al., 2016; Vekaria et al., 2017). It is thus often argued that in different contexts or in different individuals, superficially similar altruistic actions can be guided by fundamentally distinct motives (such as personal moral norms, responsibility, or empathy), which may be controlled by fundamentally distinct types of neurocognitive mechanisms (Hein et al., 2016; Piliavin and Charng, 1990; Zaki and Mitchell, 2011).

Previous studies in psychology, economics, and neuroscience have attempted to clarify this question by examining psychological motives (and their neural basis) that may underlie wealth distribution behaviors. One factor reliably influencing altruism during wealth distribution is the inequality in resources held by the actor and the recipient of a possible distribution: People are more willing to share if they possess more than the recipient (advantageous inequality, ADV) than if they possess less (disadvantageous inequality; DIS) (Charness and Rabin, 2002; Fehr and Schmidt, 1999; Gao et al., 2018; Güroğlu et al., 2014; Morishima et al., 2012; Tricomi et al., 2010). Although this consistent effect has been formalized with the same utility model across contexts, this model needs to comprise two distinct latent parameters quantifying altruism in the two contexts (i.e., decision weights on others’ interest that are specific for ADV and DIS), and these are often uncorrelated and differ strongly from each other (Gao et al., 2018; Morishima et al., 2012). These observations, together with distinct psychological accounts for the distribution behaviors in different contexts (i.e., “guilt” in the advantageous and “envy” in the disadvantageous inequality context), imply that altruistic choices in the two contexts may be driven by fundamentally different processes (Fehr and Schmidt, 1999; Gao et al., 2018). Moreover, modelling studies often reveal considerable individual differences, since these altruism parameters vary strongly between different people for the same choice set (Fehr and Schmidt, 1999). While these behavioral findings confirm clear contextual and individual differences in altruism, they leave it unclear what neurocognitive mechanisms these differences arise from. Is altruistic choice taken in a fundamentally different manner from initial perceptual processing via value computations to final integrative choice mechanisms in these different contexts and by different individuals (as suggested by (Gao et al., 2018; Tricomi et al., 2010))? Or do people perceive and attend to the choice-relevant information in a different manner before employing the same decision mechanism devoted to all types of altruistic choices (as suggested by (Yu et al., 2014))? Answering these questions would help us understand the biological origins of altruism, reveal why people differ strongly in this behavioral tendency, and develop more efficient strategies to facilitate altruism.

Empirical studies of this issue may harness the fact that altruistic decisions – like all choices - are typically guided by processes unfolding at different temporal stages (Seo and Lee, 2012; Shin et al., 2021; Tump et al., 2020), including (1) initial sensory perception of the objective information relevant for assessing wealth distribution (e.g., numbers) (Nieder, 2016; Pinel et al., 2004), (2) biased representations of the subjectively decision-relevant information attributes, such as attention-guided weighing of self- vs other-payoffs (Chen and Krajbich, 2018; Teoh et al., 2020), (3) integration of all these attributes and subjective preferences into decision values (Collins and Frank, 2018; Harris et al., 2018; Hutcherson et al., 2015), and (4) decision processes that transform the decision values into motor responses (O’Connell et al., 2012; Polanía et al., 2014). However, functional neuroimaging research has mostly employed fMRI to identify spatial patterns of neural activity that underlie wealth distribution behaviors in different inequality contexts, without explicitly examining the timecourse of information processing (Gao et al., 2018; Hu et al., 2015; Yu et al., 2014). This has revealed ambiguous results. On the one hand, distribution behavior in both contexts correlates with activity in brain regions commonly associated with motivation (e.g., the putamen and orbitofrontal cortex); on the other hand, either context also leads to activity in a set of distinct areas (the dorsolateral and dorsomedial prefrontal cortex in advantageous and the amygdala and anterior cingulate cortex in disadvantageous inequality) (Gao et al., 2018; Yu et al., 2014). Moreover, neuroanatomical research shows that for advantageous inequality only, individual variations in altruistic preferences relate to gray matter volume in the temporoparietal junction (TPJ) (Morishima et al., 2012). While these studies thus suggest that different brain areas may be activated in different contexts, it is unclear whether this indeed reflects the involvement of different hard-wired neural choice mechanisms or reflects the differences in cognitive processing during certain stages.

Moreover, recent studies combined computational modelling with fMRI techniques to show that the value of altruistic choice can be modelled as the weighted sum of self- and other-interest, and that different attributes are integrated into an overall value signal correlating with BOLD activity in ventromedial prefrontal cortex (vmPFC) (Crockett et al., 2017, 2013; Hutcherson et al., 2015). However, since these studies neither formally examined the difference in altruistic choices between advantageous and disadvantageous inequality contexts, nor provided temporally precise evidence for the decision mechanisms of altruistic choice, they can hardly address the questions whether and how different mechanisms are involved in altruism across different temporal processing stages under different contexts (Crockett et al., 2013, 2008; Gao et al., 2018). In principle, the available results may be interpreted to reflect either distinct perceptual or attentional biases to different choice-relevant information (Swart et al., 2018; Teoh et al., 2020), or shifts in decision weights for certain attributes (e.g., self-payoff) during value integration (Crockett et al., 2013; Gao et al., 2018; Saez et al., 2015), or different neural mechanisms that link decision values to motor responses (Polanía et al., 2014). Distinguishing these alternative scenarios would require use of an experimental and modelling framework that allows formal separation of the initial perceptual versus the subsequent valuation and decision processes involved in distribution behavior, as well as neural measurement techniques that can identify such processes at the level of temporally precise neural dynamics.

In the current study, we take such an approach. We combined a modified dictator game that independently varies payoffs to a player versus another person, and thereby also the inequality between both players, with electroencephalography (EEG) and sequential sampling modelling (SSM). This allowed us to identify electrophysiological markers of the perceptual processing of the decision-relevant information (i.e., stimulus-locked ERPs related to the payoffs and the resulting inequality) as well as of the processes integrating this information into a decision variable used to guide choice (i.e., response-locked evidence accumulation (EA) signals) (Balsdon et al., 2021; Hutcherson et al., 2015; Krajbich et al., 2015; Nassar et al., 2019). Thus, our approach differs from that of fMRI studies identifying brain areas involved in valuing of own and others’ payoffs (Fehr and Schmidt, 1999; Morishima et al., 2012; Saez et al., 2015), since the temporal resolution of fMRI measures makes it very difficult to separate response-locked decision making processes from stimulus-locked perceptual processes, and to examine the independent dynamics of these processes during distribution decisions.

Our approach is also motivated by studies of nonsocial decisions showing that SSMs may provide a useful framework for investigating the temporal dynamics of the processes that integrate different choice attributes into the decision outcome (Harris et al., 2018; Maier et al., 2020). Many studies have shown that SSMs can identify these processes not just computationally, but also at the neural level, for both perceptual (Brunton et al., 2013; Kelly and O’Connell, 2013; Ossmy et al., 2013) and value-based decision making (Glaze et al., 2015; Hutcherson et al., 2015; Pisauro et al., 2017; Polanía et al., 2014). The SSM framework provides a formal way to predict the temporal dynamics of processes that integrate evidence for one choice option over another for the temporal period leading up until choice, and to separate these from initial perceptual processes time locked to stimulus presentation. Neural signals corresponding to these predicted evidence-accumulation signals have been identified with EEG for perceptual decision making across different sensory modalities or stimulus features (Kelly and O’Connell, 2013; O’Connell et al., 2012; Wyart et al., 2012) as well as for value-based decision making (Pisauro et al., 2017; Polanía et al., 2014). These studies have identified evidence accumulation processes either as the model-free build-up rate of the centroparietal positivity (CPP) (Kelly and O’Connell, 2013; Loughnane et al., 2018, 2016; O’Connell et al., 2012) or in SSM-prediction-based neural signals measured over parietal and/or frontal regions (Pisauro et al., 2017; Polanía et al., 2014). Both types of neural signals are commonly interpreted as reflecting integration of the choice-relevant evidence to reach a decision, rather than basic motor planning which is usually identified by a fundamentally different neural signal, the contralateral action readiness potential (Kornhuber and Deecke, 2016; Schurger et al., 2021). The cortical origins of these signals may in principle correspond to locations identified by fMRI studies of corresponding SSM-predicted evidence accumulation traces, but note that these latter studies cannot study the temporal dynamics of these signals and unambiguously separate them into stimulus-locked perceptual versus response-locked decision processes (Gluth et al., 2012; Hare et al., 2011; Hutcherson et al., 2015; Rodriguez et al., 2015).

Here we apply this approach and use SSMs fitted to individuals’ wealth distribution behaviors to predict the underlying neural evidence accumulation dynamics. We then employ these predicted EA signals in our EEG analyses to examine whether a similar neural choice system accumulates the choice-relevant evidence in both inequality contexts, or whether distinct neural systems implement this decision process for the different contexts. Then, we examined whether the different features of each choice problem that ultimately need to be integrated in the choice-relevant evidence – that is, the specific payoffs available to oneself and the other person, as well as the inequality context – are initially processed in a different manner for different contexts and in different individuals. This allowed us to directly approach the question whether contextual and individual differences in altruism arise from the use of fundamentally different mechanisms across different cognitive processing stages, or rather from differences in initial processing of the choice-relevant information that is ultimately integrated into the same valuation and decision mechanisms.

## Results

We recorded 128-channel EEG data from healthy participants playing a modified Dictator Game (DG). On each trial of this task, participants played as proposers and chose between two possible allocations of monetary tokens between themselves and an unknown partner. We systematically varied the allocation options from trial to trial so that in half of the trials, participants received less than their partners for both choice options (disadvantageous context (DIS)) and in the other half they got more than their partners for both options (advantageous context (ADV)). These two types of trials were randomly intermixed and were only defined by the size of the payoffs presented on the screen.

On each trial, we presented the two options sequentially, to allow clear identification of timepoints at which the information associated with each option was processed (Figure 1A, see SI Materials and Methods for details). This sequential presentation allowed us to establish the inequality context with the presentation of the first option, without having to explicitly instruct participants about the different contexts. We then studied individuals’ sensitivity to self-payoff, other-payoff, and inequality level by focusing on how choices of the second option depended on the change in these variables from the first to the second option. Importantly, as shown in the payoff schedule of all trials (Fig. S1), we matched self-/other-payoff differences and the resulting absolute levels of inequality across both contexts and also across the second and the first options (Fig. S1 middle and right panels). This allowed us to compare choices and response times, model-defined neural choice processes timelocked to the response, and neural processing of different stimulus information (self- and other-payoff, inequality) between the two contexts.

**Figure 1.**
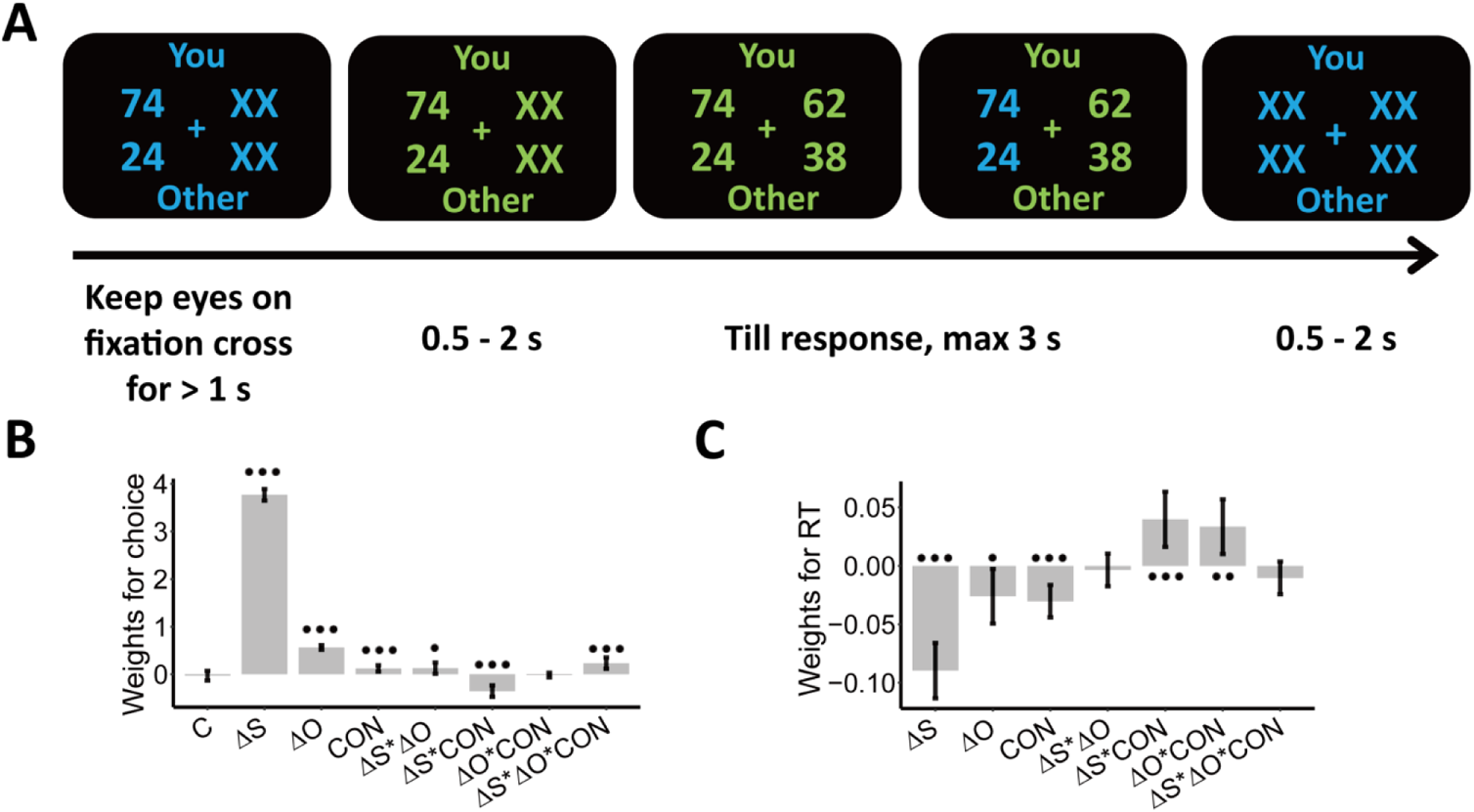
Experimental design and behavioral results. We employed a modified dictator game (DG) to measure individuals’ wealth distribution behaviors. (A) Example of display in a single trial. In the task, participants played as proposers to allocate a certain amount of monetary tokens between themselves and anonymous partners. In the beginning of each trial, participants were presented with one reference option in blue and were asked to keep eyes on the central cross for at least 1 s to start the trial, as indicated by the change in font color from blue to green. When the 2^nd^ option was presented, participants had to choose between the two options within 3 s. The selected option was highlighted in blue before the inter-trial interval. Font color assignment to phases (i.e., blue and green to response) was counterbalanced across participants. (B) Payoff information and context affect choice systematically. Generalized linear mixed-effects model shows the effects of multiple predictors on probability to choose the 2^nd^ option; (C) Payoff information and context affect reaction times systematically. Linear mixed-effects model shows the effects of multiple predictors on response times (RTs). ΔS, Self-payoff Change; ΔO, Other-payoff Change; CON, Context; ···, *p* < 0.001; ··, *p* < 0.01; •, *p* < 0.05.

Based on the model fits and their predicted response-locked evidence accumulation EEG traces, we first tested whether similar or different neural processes (i.e., brain regions or physiological markers) underlie the ultimate choice process in the two inequality contexts, in similarity to how this has been studied for other types of decisions (Polanía et al., 2014). Then, we clarified whether neural processing of the stimulus information – which subsequently feeds into the decision processes - differs across contexts and individuals. For this analysis, we examined stimulus-locked event-related potentials (ERPs), in a way that has also been used to differentiate neural processing of decision-relevant features in non-social value-based decision making (e.g., perceptions of health and taste of food items) (Harris et al., 2018). Finally, we explored how individual differences in altruism are related to large-scale information communications between regions associated with these two sets of processes (i.e., response-locked decision processes and stimulus-locked perceptual processes), by examining inter-regional synchronization in the gamma-band frequency (30-90 Hz). This last analysis was motivated by the consideration that evidence accumulation processes need to integrate evidence input from different neural sources (e.g., perceptual processes) (Polanía et al., 2014), and by the proposal that coherent phase-coupling in gamma band between different groups of neurons may serve as a fundamental process of neural communication for information transmission (Bosman et al., 2014; Fries, 2009, 2005; Vinck et al., 2013), as already shown for non-social value-based decisions (Polanía et al., 2014; Siegel et al., 2008).

### Behavior: Altruism depends differentially on self- and other-payoffs across contexts

Before performing model-based analyses, we ran model-free linear mixed-effects regressions to establish that the choice-relevant information [i.e., self-payoff, other-payoff, and inequality context (ADV and DIS)] indeed systematically affects individual wealth distribution choices. These analyses confirmed that both self-payoff and other-payoff were important factors underlying individuals’ choice.

Specifically, participants chose the second option more often when either they or the receiver profited more from this choice (main effect Self-payoff Change (ΔS): beta = 3.77, 95% CI [3.65 – 3.89], *p* < .001; main effect Other-payoff Change (ΔO): beta = 0.56, 95% CI [0.51 – 0.61], *p* < .001, ΔS(ΔO): participants’ own (partners’) payoff change between the 2^nd^ and the 1^st^ option) (Table S1, Figure 1B). However, participants were less influenced by changes in their own payoff when they had more money than the other (ADV, interaction Self-payoff Change (ΔS) and Context (CON): beta = −0.35, 95% CI [−0.47– −0.23], *p* < .001) or when the receiver got lower payoffs from this choice (decreasing other-payoff, interaction Self-payoff Change (ΔS) and Other-payoff Change (ΔO): beta = 0.13, 95% CI [0.01– 0.24], *p* = .03). This latter effect was particularly notable when the participants had more money than the receiver (ADV; three-way interaction Self-payoff Change (ΔS), Other-payoff Change (ΔO), and Context (CON): beta = 0.23, 95% CI [0.11 – 0.35], p < .001; Table S1, Figure 1B). For visualizations of these effects, please see SI results and Fig. S2A. Please note that we also constructed simpler models without interaction effects and/or main effects, but model comparison analyses favored the full model (Table S1). An additional linear mixed-effects regression model suggested that the presentation order (i.e., 1^st^ or 2^nd^) of options would not affect individuals’ equal/unequal choices (see SI Results and Table S2).

Self-payoff, other-payoff, and context also jointly affected how quickly participants took their decisions. Choices were faster for larger absolute values of self-payoff change (main effect Self-payoff Change (|ΔS|): beta = −0.09, 95% CI [−0.11 – −0.07], *p* < .001) and other-payoff change (main effect of Other-payoff Change (|ΔO|): beta = −0.03, 95% CI [−0.05 – −0.003], *p* = .03) (Figure 1C). Again, both these effects were different for the two inequality contexts, with reaction times more strongly affected in the disadvantageous inequality context (interaction between Self-payoff Change (|ΔS|) and Context (CON): beta = 0.04, 95% CI [0.02 – 0.06], *p* < .001; interaction between Other-payoff Change (|ΔO|) and Context (CON): beta = 0.03, 95% CI [0.01 – 0.06], *p* = .005; Table S3, Figure 1C). These effects are consistent with the central assumption of the SSM framework that stronger (weaker) evidence will speed up (slow down) evidence accumulation and resulting choice, thereby already suggesting that an SSM-based decision process may integrate self- and other-payoff to guide individual decisions (For visualizations of these effects, see Fig. S2B).

### Model-based EEG reveals similar parietal evidence accumulation across contexts

To address the question whether distribution choices were supported by a similar or different neural decision processes across both inequality contexts, we fitted a dynamical sequential sampling model (SSM) to participants’ behavioral data and used it to predict neural evidence accumulation (EA) signals for the two contexts. Please see SI Materials and Methods for comparison of alternative models and the details of the best-fitting model we used for our analysis.

Our analyses revealed comparable SSM-based EA signals over similar parietal regions for both contexts, and no context-specific EA signals that would indicate use of fundamentally different neural choice mechanisms in the different contexts. Specifically, we first fitted the SSM by categorizing trials as “equal” or “unequal” choices, based on whether the participant selected the option with more equal or less equal distribution of monetary tokens between both players. For each trial, the model used the subjective value difference (VD) between the more equal option and the more unequal option (computed using the Charness-Rabin utility model, see SI Materials and Methods) as its input to predict moment-by-moment evidence accumulation signals until the timepoint when the decision was made. For this, we used the Ornstein-Uhlenbeck choice model (OU), which assumes a leaky accumulation-to-bound process (Bogacz et al., 2006) and was the model favoured by our model comparisons (see SI Materials and Methods). This model allowed us to fit several free parameters corresponding to different aspects of preference and the choice process: the relative decision weight on others’ payoffs *ω* (altruistic preference), decision threshold *α* (response caution), starting point *β* (response bias), and drift rate modulator *κ* (sensitivity to equality-related information), as well as the more technical parameters leak strength *λ* and non-decision time *τ* (see SI Materials and Methods for a detailed model description). By including these parameters, we could examine the effects of context on both basic altruistic preference (i.e., *ω*) and the decision process that integrates the resulting subjective values into the distribution choice (i.e., *α*, *β*, and *κ*).

To simulate the evidence accumulation process, we averaged 500 EA traces generated by the participant-specific fitted model for the given context and the payoffs on each trial. Model simulations showed that these traces were good approximations of the EA processes underlying choice, since the fitted model could both capture choices and RTs across the two contexts. For both types of choices (equal/unequal) and contexts (ADV and DIS), the sensitivity/specificity of the data simulated by the model was higher than 83% (Figure 2A left panel) and the balanced accuracy was higher than 89% (Figure 2A right panel, SI Materials and Methods). The model also correctly captured response speed effects, with correct predictions that choices are faster during advantageous inequality overall (RT in ADV: 0.80 ± 0.04 s, DIS: 0.84 ± 0.04 s, ADV vs. DIS: 95% CI [−0.06, −0.02], Cohen’s d = −0.83, *t*(37) = −5.12, *p* < 0.001), and faster for equal (unequal) choices in DIS (ADV) (DIS: RT equal choice: 0.78 ± 0.03 s, unequal choice: 0.91 ± 0.04 s; equal vs unequal choice: 95% CI [−0.16, −0.10], Cohen’s d = −1.51, *t*(37) = −9.31, *p* < 0.001; ADV: RT unequal choice: 0.73 ± 0.03 s, equal choice: 0.87 ± 0.04 s; unequal vs equal choice: 95% CI [−0.17, −0.11], Cohen’s d = −1.54, *t*(37) = −9.47, *p* < 0.001, Figure 2B).

**Figure 2.**
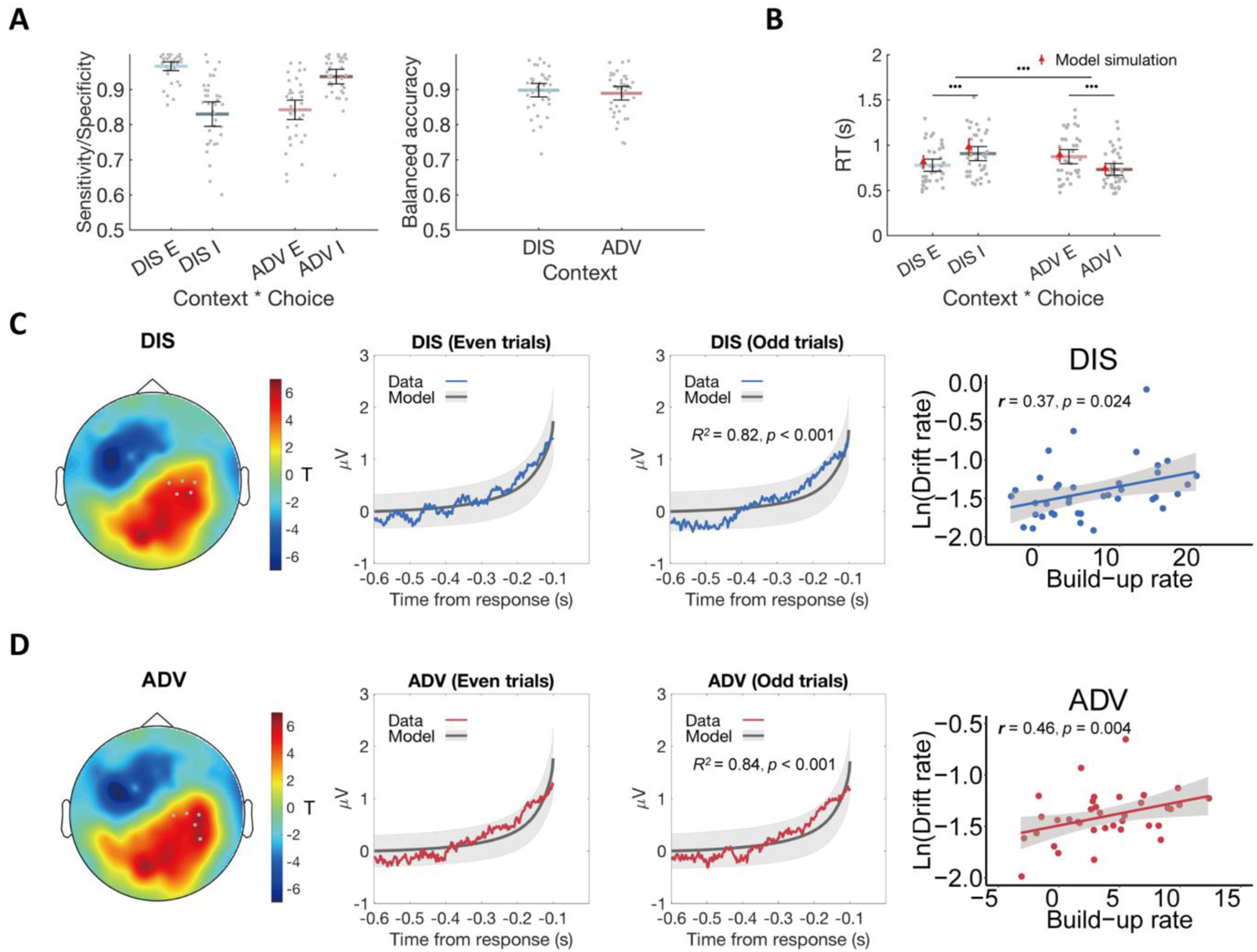
Model fits and relationship between ERP signal and model predictions. (A) The OU model predicts choices across contexts. Left panel: Model performance on Sensitivity of equal choice (E) /Specificity of unequal choice (I) in each context. Right panel: Model performance on balanced accuracy in each context. DIS, disadvantageous context; ADV, advantageous context; E, equal choice; I, unequal choice. (B) The OU model recovers RT effects over context and choice in participants’ behavioral data. Black error bars display means ± 95% confidence intervals (CIs). Each grey dot indicates one participant. The red triangle dots and error bars represent model simulation mean and 95% CIs. ···, *p* < 0.001. (C) & (D) Comparable neural evidence accumulation signals in both contexts. For each context (C for DIS, D for ADV), response-locked epochs were divided into even and odd trials to perform cross-validation analyses. Leftmost panels show the topographic scalp distributions of associations between ERP amplitude and OU model predictions in even trials. Grey dots highlight channels that survived the threshold. Middle-left panels show the observed averaged ERP data in the identified clusters shown in leftmost panels (colored lines) and normalized OU model predictions for even-numbered trials. Middle-right panels show the relationships between the model predictions and ERP signals of the independent half of data (odd-numbered trials) in the identified clusters shown in leftmost panels. Colored lines represent average ERP data extracted from the identified clusters. Grey lines represent the means of model-predicted EA traces, and grey shaded areas represent ± 1 standard errors of the mean (SEM) of the model-predicted EA traces. Rightmost panels show the correlations between the built-up rates of the identified clusters and the OU model parameter of drift rates across participants.

To identify the corresponding neural traces, we performed cross-validated regressions of EEG signals on the SSM-based EA predictions. We divided trials in each context into even- and odd-numbered trials and used the former to identify channels in which ERP magnitude correlated with the model-predicted EA traces. We then used regressions to test the model predictions for the data extracted from these channels for the independent odd-numbered trials (see SI Materials and Methods for details). In both contexts, we observed model-predicted neural EA signals in the same channels located over parietal regions (Figure 2C & 2D leftmost and middle-left panels, *p_Bonferroni_* < 0.05), and ERP waveforms extracted from these channels for the independent odd-numbered trials showed a significant correspondence with the model-predicted EA signals (DIS: *R^2^* = 0.82, *p* < 0.001; ADV: *R^2^* = 0.84, *p* < 0.001, Figure 2C & 2D middle-right panel). These neural expressions of EA signals did not differ between advantageous and disadvantageous inequality contexts, as ascertained by in-depth comparisons of the observed ERP data (t-tests for each EEG electrode did not reveal any significant differences, see SI Materials and Methods for detailed descriptions of statistics).

We also confirmed that decision signals were comparable across contexts using an alternative model-free analysis approach established in the literature (Kelly and O’Connell, 2013). This analysis calculates the build-up rates of the response-locked ERP waveforms in the clusters shown in Figure 2C-D and correlates this rate with the drift rate modulator derived from the fitted SSM for each context. ERP build-up rates were indeed associated with the drift rates in both inequality contexts (DIS: *r*(37) = 0.37, *p* = 0.024; ADV: *r*(37) = 0.46, *p* = 0.004) (Figures 2C & 2D rightmost panels), also supporting the conclusion that similar brain areas implement evidence-accumulation processes for both inequality contexts. Moreover, the convergence of the results from both analysis approaches highlights the similarity of the signals observed here to those observed in the same parietal sensors for other types of decision making, such as perceptual and non-social value-based choices (Pisauro et al., 2017; Polanía et al., 2014).

### Contextual differences in aspects of unified decision mechanism

Given the EEG evidence for a comparable neural decision mechanism across both inequality contexts, we next examined which specific aspects of this mechanism may vary to produce the behavioral differences the two inequality contexts. We focused on four decision-relevant parameters, including the relative decision weight *ω* capturing perception of other-payoff versus self-payoff (i.e., altruistic preferences), the decision threshold *α* indexing response caution, the starting point *β* quantifying response bias, and the drift rate *κ* measuring the sensitivity of evidence integration (see Fig. S3 for the distributions of response urgency with increasing time (leak strength *λ*) and speed of pre-decision information processing (non-decision time (nDT) *τ*)).

The analysis revealed that only two parameters differed significantly across inequality contexts. In the advantageous context, perception was characterized by a higher weight on others’ payoffs (*ω*) (ADV: 0.15 ± 0.02, DIS: 0.10 ± 0.02, ADV vs DIS: 95% CI [0.014, 0.10], Cohen’s d = 0.44, *t*(37) = 2.74, *p* = 0.037, Bonferroni correction), but the decision mechanism showed a lower decision threshold (*α*) (ADV: 1.11 ± 0.06, DIS: 1.34 ± 0.09, ADV vs DIS: 95% CI [−0.33, −0.13], Cohen’s d = −0.77, *t*(37) = −4.78, *p* < 0.001, Bonferroni correction, Figure 3A). These findings indicate that when individuals are initially endowed with more money than the other, they require less evidence to take a choice, but crucially consider the others’ payoffs more strongly.

**Figure 3.**
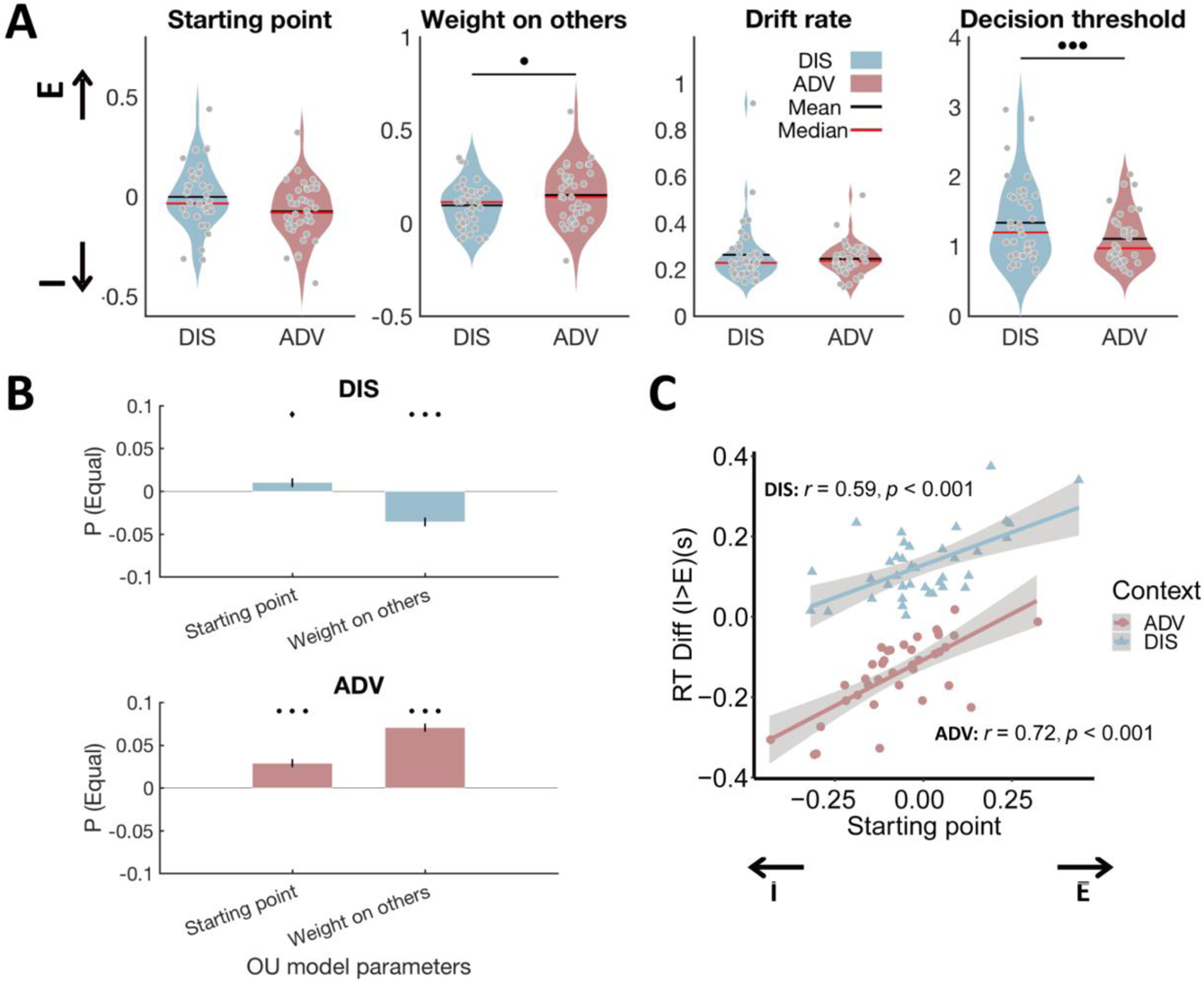
Model parameters differ between inequality contexts and show the expected relations to model-free behavioral data. (A) Relative to DIS, ADV increased weight on others, and reduced decision threshold. Each grey dot represents one participant. E, starting point closer to equal option; I, starting point closer to unequal option. (B) Multiple regressions show that the probability to choose the equal option related to a starting point closer to the equal option in both contexts, and a higher weight on others in ADV and a lower weight on others in DIS context. (C) Individuals with starting point closer to the equal option responded more slowly for unequal relative to equal choices in both contexts. E, starting point closer to equal option; I, starting point closer to unequal option.···, *p* < 0.001; ··, *p* < 0.01; ♦, *p* < 0.1.

Reassuringly, these differences in model parameters captured the key model-free behavioral differences between contexts: The individual probability to choose the equal option was significantly correlated with the weight on other-payoff (regression, ADV: *beta*(35) = 0.07, *SE* = 0.005, *p* < 0.001; DIS: *beta*(35) = −0.04, *SE* = 0.005, *p* < 0.001) and the bias in starting point (DIS: *beta*(35) = 0.01, *SE* = 0.005, *p* = 0.064; ADV: *beta*(35) = 0.03, *SE* = 0.005, *p* < 0.001, Figure 3B). Similarly, individuals with a stronger bias in starting point towards the equal option took faster equal decisions and slower unequal decisions (Pearson correlation between starting point and difference in reaction time between unequal choice and equal choice in DIS: *r*(37) = 0.59, *p* < 0.001; and in ADV: *r* (37) = 0.72, *p* < 0.001, Figure 3C). Irrespective of these differences between contexts, correlation analyses again suggested that people may employ a comparable overall decision mechanism across the two contexts, since all model parameters (Fig. S4, minimum correlation: *r* = 0.30, *p* = 0.06; maximum correlation: *r* = 0.90, *p* < 0.001) and the probability to choose the equal option (Fig. S5; *r* = −0.62, *p* < 0.001) were highly correlated across the two inequality contexts.

Together with the ERP data, these findings imply that even though participants took their choices in both contexts using a similar decision mechanism, various aspects of this mechanism (e.g., decision thresholds) and of the processes supplying the choice-relevant information (e.g., weights on other-payoff) differ across individuals and contexts. These behavioral and modelling findings motivated us to examine how exactly the *neural* processes extracting choice-relevant information form the stimuli may differ between the two inequality contexts.

### Contextual differences in altruistic choice reflect different neural processing of self-payoffs

To investigate neural processing of the choice-relevant information contained in the stimuli, we examined stimulus-locked ERP signals with general linear models (GLMs). Since the payoff options were presented sequentially, we assumed that people initially evaluate the changes (from first to second option) in their own or others’ payoffs to then integrate this information into the choice process. We thus switched from a choice-centered analysis approach (i.e., SSM and response-locked EEG analyses discussed above) to a stimulus-centered approach and analyzed how stimulus-locked ERPs reflected different aspects of the payoff information on the screen. To do so, we entered stimulus-locked ERP signals into a linear regression model (see SI for details) in which participants’ own (partners’) payoff change between the 2^nd^ and 1^st^ option [Δ*S* (Δ*O*)] were included as predictors. This linear regression produced a set of estimated coefficients measuring the parametric effect strengths of Δ*S* and Δ*O* on neural activity for each channel and time window in each context and participant.

In these GLM analyses, we focused on the effect size measures to quantify with permutation tests corrected for multiple comparisons (see SI Materials and Methods for details) when and where EEG signals reflected self-payoff changes (Δ*S*) and other-payoff changes (Δ*O*). For the significant clusters emerging from these analyses, we then conducted post-hoc analyses of raw ERP amplitudes to characterize the identified effects in more detail. This analysis approach allowed us to test for two different possible scenarios of how context-effects in distribution behavior may relate to neural perception processes.

On the one hand, while participants may always guide their choice by both the self-payoff change (Δ*S*) and the other-payoff change (Δ*O*), they may focus more strongly, or at different timepoints, on their own payoff or the receiver’s payoff in the two contexts (Scenario 1) (Harris et al., 2018). On the other hand, participants may guide their choice more by fairness considerations (i.e., inequality estimates) that are at conflict with the motivations to maximize either self-payoffs or other-payoffs in the two contexts (*12, 14*). If this scenario holds, then ERPs should relate to inequality levels in opposite ways for the two contexts (Scenario 2), since a more unequal second option relates to increased self-payoff and/or decreased other-payoff in ADV, but to decreased self-payoff and/or increased other-payoff in DIS.

Our results support scenario 2: A conjunction analysis revealed no significant ERP component that showed similar effects of self-payoff change across the two contexts. Instead, when contrasting the whole-brain *β*_1_ maps between the two contexts, we identified two different spatial-temporal clusters that showed opposite effects of self-payoff changes between contexts (i.e., earlier effect: from ~ 240 to 360 ms after stimulus onset, *p_cluster_* = 0.04, Figure 4A; later effect: from ~ 440 to 800 ms after stimulus onset, *p_cluster_* < 0.001, Figure 4C). In the earlier time window, more negative centroparietal neural responses were associated with self-payoff increases (positive Δ*S*) in the ADV context (−0.09 ± 0.04, Mean *β*_1_ ± SE, 95% CI [−0.17, −0.01], Cohen’s d = −0.36, *t*(37) = −2.23, *p* = 0.03) but with self-payoff decreases (negative Δ*S*) in the DIS context (0.08 ± 0.03, 95% CI [0.01, 0.15], Cohen’s d = 0.40, *t*(37) = 2.45, *p* = 0.02; ADV vs. DIS: 95% CI [−0.28, −0.06], *t*(37) = −3.21, Cohen’s d = −0.52, *p* = 0.003, Figure 4A right panel). In the later time window (~ 440 to 800 ms after stimulus onset), however, more *positive* centroparietal neural responses related to self-payoff increases in the ADV context (0.10 ± 0.03, 95% CI [0.04, 0.16], Cohen’s d = 0.52, *t*(37) = 3.21, *p* = 0.003) but to self-payoff decreases in the DIS context (−0.12 ± 0.03, 95% CI [−0.18, −0.05], Cohen’s d = −0.60, *t*(37) = −3.68, *p* < 0.001; ADV vs. DIS: 95% CI [0.11, 0.32], *t*(37) = 4.30, Cohen’s d = 0.70, *p* < 0.001, Figure 4C right panel). Thus, these ERP effects more likely relate to inequality-conflict processing (scenario 2) than to common perceptions of self-payoff change across different contexts (scenario 1).

**Figure 4.**
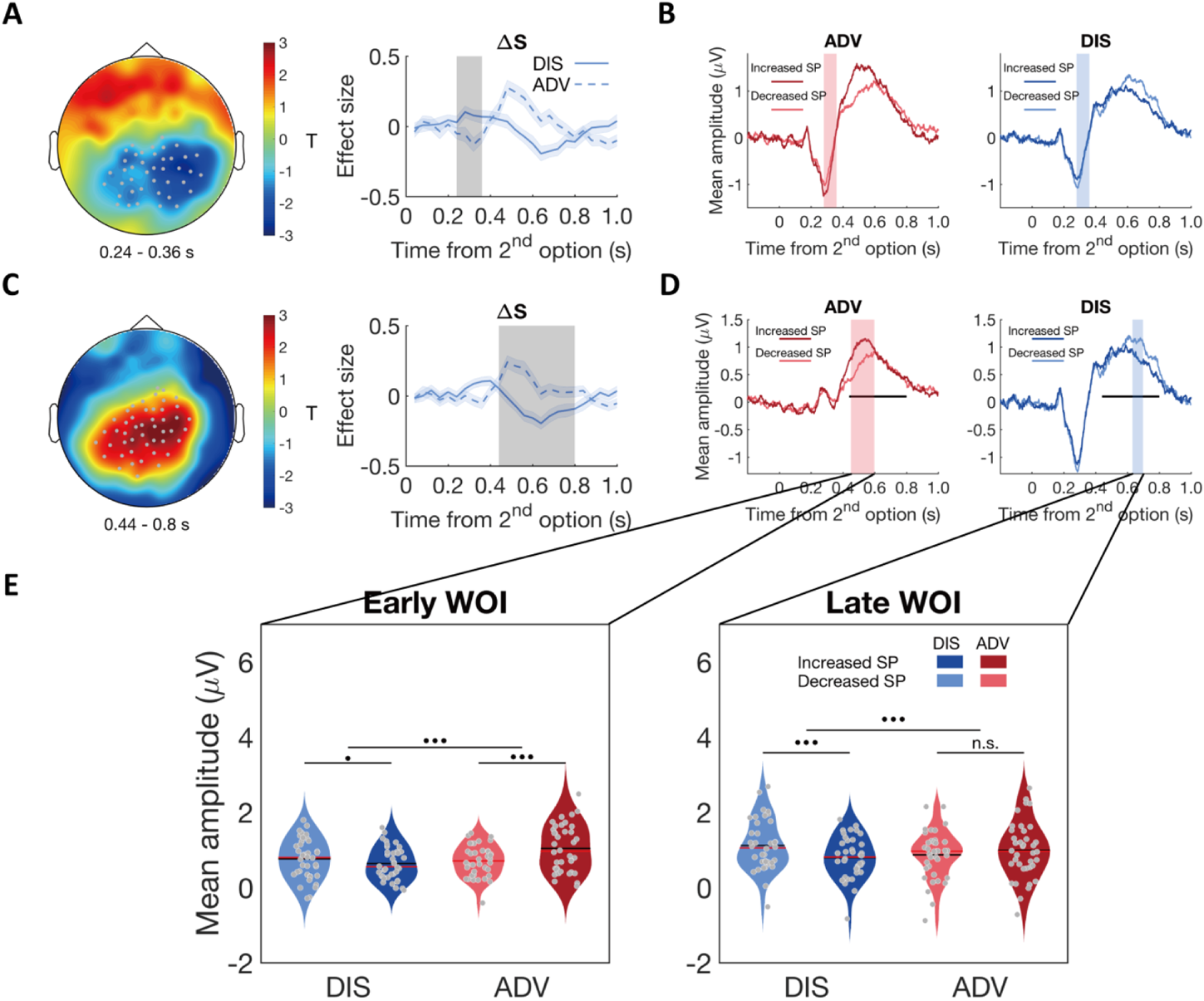
Contextual differences in neural inequality processing revealed by stimulus-locked ERPs. (A) & (C) ERP correlations with self-payoff change (Δ*S*) show opposite signs for the two inequality contexts, in the time windows of ~240 – 360 ms (A) and ~440 – 800 ms (C): left panels, topographic scalp distributions of context differences; right panels, temporal dynamics of the parametric effect strengths of self-payoff change (Δ*S*) in the identified clusters. Grey dots highlight channels which survived the threshold, grey shaded areas indicate the durations of the significant effects, and colored shaded areas indicate ± 1 SEM in (A) and (C). (B) and (D) Average ERP waveforms for the parametric effect of Δ*S* in each context during the windows of ~240 – 360 ms (B) and ~440 – 800 ms (D) (For statistic information, see SI Results). Significant clusters surviving the cluster correction for multiple comparisons at *p* < 0.05 are reported. Increased SP, trials in which the 2^nd^ option increase self-payoff; Decreased SP, trials in which the 2^nd^ option decrease self-payoff. Colored shaded areas indicate the duration of significant interaction between context and self-payoff change in (B). Black lines indicate the duration of significant interaction between context and self-payoff change identified in (C), and colored shaded areas in (D) indicate separate durations of the significant interaction effect of self-payoff change and context over ERP waveforms in the time window identified in (C). (E) Inequality processing occurs at different time periods in the two contexts. For the significant cluster shown in (C), Δ*S* effects lasted from ~ 450 to 600 ms after stimulus onset in ADV (D, pink shaded area), and from ~ 630 to 700 ms after stimulus onset in DIS (D, blue shaded area). ERP responses are averaged magnitudes derived from each window of interest (WOI) in each condition. ERP responses to self-payoff change are stronger in an earlier WOI (~ 450 to 600 ms, left panel) in ADV context, and stronger in a later WOI (~ 630 to 700 ms, right panel) in DIS context. For the early WOI (left panel), increased self-payoff (Increased SP) was related to stronger neural responses than decreased self-payoff (Decreased SP) in ADV, but there was no difference of neural responses between decreased self-payoff and increased self-payoff in DIS (left panel). For the late WOI, decreased self-payoff (Decreased SP) was related to stronger neural responses than increased self-payoff (Increased SP) in DIS, but there was no difference of neural responses between increased self-payoff and decreased self-payoff in ADV (right panel). For statistic information, see SI Results. Increased SP, trials in which the 2^nd^ option increase self-payoff; Decreased SP, trials in which the 2^nd^ option decrease self-payoff. Red lines, Median; black lines, mean. ···, *p* < 0.001; •, *p* < 0.05; n.s., not significant.

Interestingly, context effects in inequality-related neural processing were mainly evident in ERP correlations with self-payoff. The same analyses of *β*_2_ maps revealed no ERP components related to other-payoff changes (Δ*O*), neither across contexts or differing between contexts. We also examined temporal dynamics of the parametric effect strengths of other-payoff change (Δ*O*) in the specific clusters identified for the context-dependent effects of self-payoff change (Δ*S*) in the above analyses, but found no significant effects (Fig. S6).

Post-hoc analyses of the ERP components corresponding to the effects identified in the above regression analyses also confirmed scenario 2. For this analysis, we categorized trials in each context into trials in which the second option increases self-payoff (i.e., more equal in DIS and more unequal in ADV) or decreases self-payoff (i.e., more unequal in DIS and more equal in ADV). For both time windows, ERPs associated with the increase and decrease of self-payoff yielded opposite neural effects in the two inequality contexts (see Figure 4B & 4D; for stats see SI Results). Importantly, a spatially-unbiased interaction analysis between context (ADV vs DIS) and self-payoff change (Increased SP vs Decreased SP) in either time window revealed that people processed self-payoff-related inequality earlier in the advantageous context (Figure 4D and 4E, for statistic information, see SI Results and SI Materials and Methods; for a temporally agnostic ERP analysis confirming these results, see Fig. S7). These timing differences suggest that the different behavior in the two inequality contexts may reflect differences in how and when participants focus neural processing on the inequality resulting from the choice options, and that this assessment is mainly based on differences in self-payoffs.

### Individual differences in altruistic choice relate to differential neural processing of other-payoffs

Choice behavior did not just differ between the two inequality contexts but also across individuals. To investigate the neural processes underlying these individual differences, we divided participants into more altruistic (MA) and less altruistic (LA) based on their weight on others’ payoff (ω, median-split) and compared ERPs indexing neural processing of self-related and other-related payoffs (using the same GLM analyses and permutation tests corrected for multiple comparisons, see SI Materials and Methods). Again, we directly investigated the two scenarios outlined above: Under scenario 1, behavioral differences may emerge from a (temporally) distinct focus on self- or other-related payoffs whereas under scenario 2, the differences may be linked to differential processing of inequality signals, which relate to payoffs in opposite ways in the two contexts.

The results of this analysis were more consistent with scenario 1, since individual differences were mainly related to variations in neural processing of other-payoffs, especially in disadvantageous inequality context. Specifically, during disadvantageous inequality, other-payoff was associated with a more negative ERP response in the less altruistic group (LA: −0.16 ± 0.06, Mean *β*_2_ ± SE, 95% CI [−0.28, −0.04], Cohen’s d = −0.64, *t*(18) = −2.78, *p* = 0.02; MA: 0.10 ± 0.04, 95% CI [0.02, 0.19], Cohen’s d = 0.56, *t*(18) = 2.46, *p* = 0.01). The effect was most strongly expressed over centrofrontal regions (*p_cluster_* = 0.03, post-hoc comparison: 95% CI [−0.43, −0.09], Cohen’s d = −0.75, *t*(18) = −3.27, *p* = 0.004, Figure 5A & 5B) and was also evident in a parametric correlation analysis between other-payoff ERP and the weights on others’ payoffs (ω) (see Fig. S8), but not for an analysis based on differences in starting point *β* (Fig. S9).

**Figure 5.**
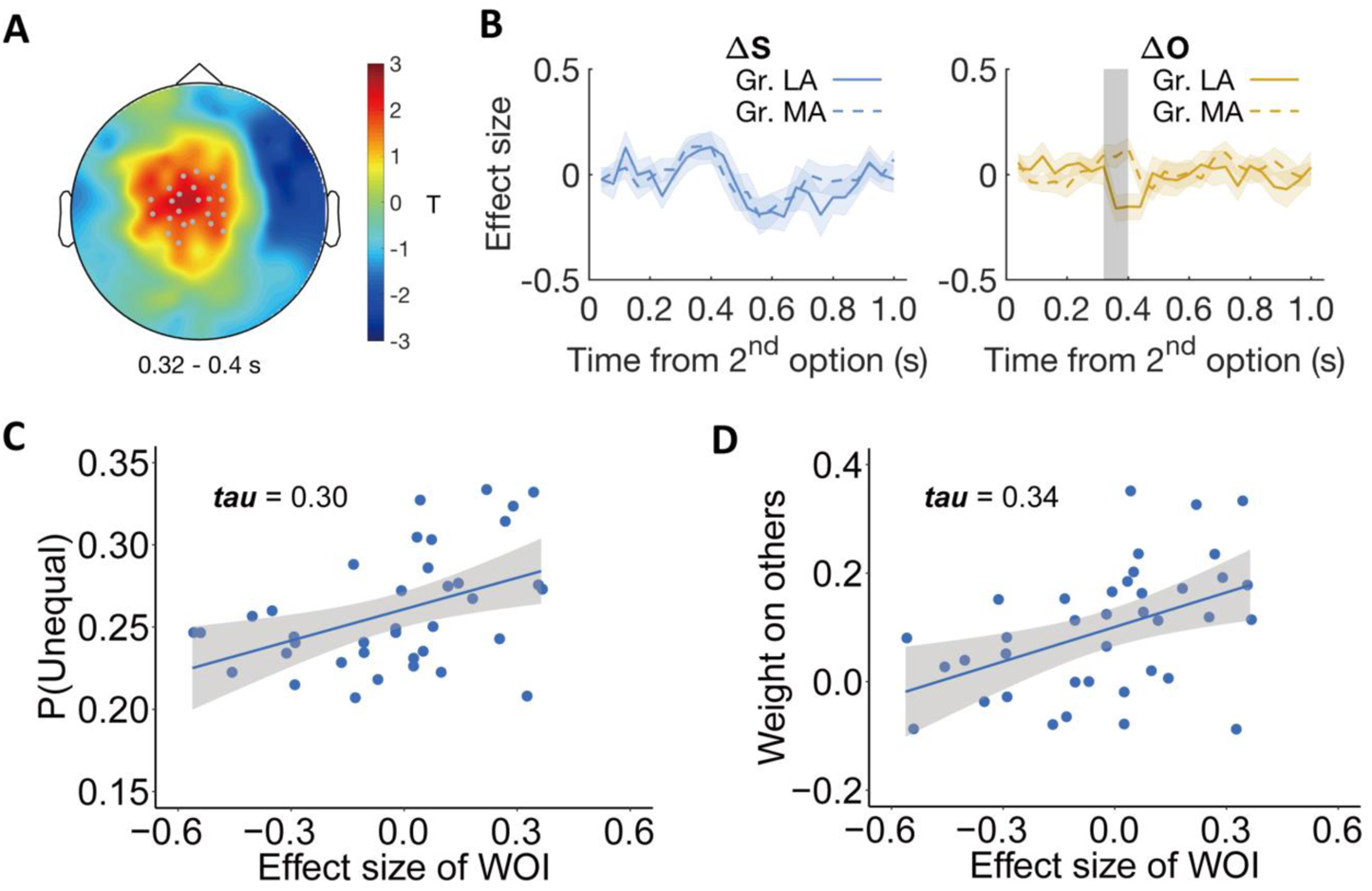
Individual differences relate to differential neural processing of other-payoffs in the stimulus-locked ERP analysis. Participants were divided into more altruistic (MA) and less altruistic groups (LA) based on the model parameter weight on others. (A) Topographic scalp distribution of the difference between MA and LA groups in ERP correlation with other-payoff difference (Δ*O*) in the DIS context. The significant effect lasted from ~320 to 440 ms after stimulus onset. Grey dots highlight channels that survived the threshold. (B) Specificity of the effect for Δ*O*. Temporal dynamics of the parametric effect strengths of self-payoff change (Δ*S*, left) and other-payoff change (Δ*O*, right) in the identified cluster. Colored shaded areas indicate ± 1 SEM. Gr., group. Grey shaded area indicates the duration of the significant effect. Significant cluster surviving the cluster correction for multiple comparisons at *p* < 0.05 are reported. (C-D) Neural effects relate to behavior. The plots show the correlations between the effect strengths of other-payoff change (Δ*O*) in the identified cluster and the probability to choose the unequal option (C) and the OU parameter of weight on others (D).

To visualize the relationship between this neural effect and both model-free and model-based measures of altruistic preferences, we extracted the individual effect size of other-payoff change (Δ*O*) in the DIS context from the identified spatio-temporal cluster and correlated it with the individual probability to choose the more unequal option and with the weight on others’ payoffs (ω) (to avoid circular inference, we only report and illustrate these correlations but do not compute corresponding p-values). This confirmed that the stronger the neural processing of the other’s payoff difference (i.e., *β*_2_), the more likely the participants were to choose the more unequal option (*Kendall’s tau* (37) = 0.30, Figure 5C). The corresponding analyses with the model parameters also confirmed positive correlations with the individuals’ weight on others (*Kendall’s tau* (37) = 0.34, Figure 5D). The correlation analyses also confirm that these effects are only evident during the DIS context, since the corresponding correlations during the ADV context were miniscule (maximum tau-value = 0.007, minimum tau-value = −0.003, Fig. S10). The same whole-brain analyses of individual differences were also performed in the ADV context, but no significant cluster was identified. For visualization of the specific origin of the neural effect identified in the individual-difference analysis of other-payoff processing, please see SI results and Fig. S11.

To confirm that these individual differences in neural processing of choice-relevant information are unrelated to the context effects described further above, we also compared the more-versus less-altruistic groups in terms of how they processed self-payoff (as analyzed in the previous section). This showed that independent of the grouping parameter (i.e., weight on others or starting point), more altruistic and less altruistic groups exhibited comparable neural processing of self-payoff differences in those identified clusters (Fig. S12). Thus, individual differences in altruistic preferences related to processing of other-payoff, and did not relate to temporal dynamics of the neural processing of inequality or self-payoff that was associated with contextual behavioral differences.

Taken together, these analyses add evidence that although distribution choices are guided by a comparable neural decision mechanism across different people and different contexts, there can be considerable contextual and individual differences in how the information fed into these processes is initially processed: While contextual differences in choice behavior mainly relate to differences in neural processing of inequality signals, individual differences in altruism relate mainly to the processing of other-related information.

### Individual differences in altruistic preferences relate to synchronization of perception and evidence-accumulation signals

In previous studies, it has been observed that evidence accumulation processes need to integrate information from brain areas processing the stimulus dimensions relevant for choice, and that this information exchange is characterized by inter-regional gamma-band coherence (Polanía et al., 2015, 2014; Siegel et al., 2008). Given that our SSM-EEG analyses identified a comparable neural evidence accumulation process in different contexts, and that our ERP analyses revealed differential processing of the trial-wise information relevant for choice, we also examined how these two sets of processes may exchange information in order to guide the decisions. Specifically, our finding that individuals’ altruistic preferences (i.e., weight on others’ payoffs) in the disadvantageous context related to neural processing of other-payoff in centrofrontal signals led us to test whether the underlying brain areas exchange this information with the parietal areas accumulating this evidence for choice.

For each participant, we computed the debiased weighted phase lag index (dWPLI, (Vinck et al., 2011)) for gamma-band coherence between these two signals in the disadvantageous context, and compared this index of phase coupling between more- and less-altruistic participants (again median-split by ω). We found that phase coupling between the centro-frontal regions (shown in Figure 5A) and the parietal regions (shown in Figure 2C) in the DIS context was indeed significantly higher for the more altruistic group (MA: 0.010 ± 0.003, 95% CI [0.004, 0.016], Cohen’s d = 0.80, *t*(18) = 3.48, *p* = 0.003; LA: group (7.75 × 10^−4^ ± 10 × 10^−4^, 95% CI [−0.001, 0.003], Cohen’s d = 0.17, *t*(18) = 0.76, *p* = 0.46, MA vs. LA: 95% CI [0.004, 0.015], Cohen’s d = 0.99, *t*(18) = 3.38, *p* = 0.003). This effect was evident in the gamma band frequency (64 – 79 Hz) for the time window ~ 520 to 460 ms before response (*p_cluster_* = 0.04, corrected for multiple comparisons, see SI Materials and Methods and Figure 6A & 6B).

**Figure 6.**
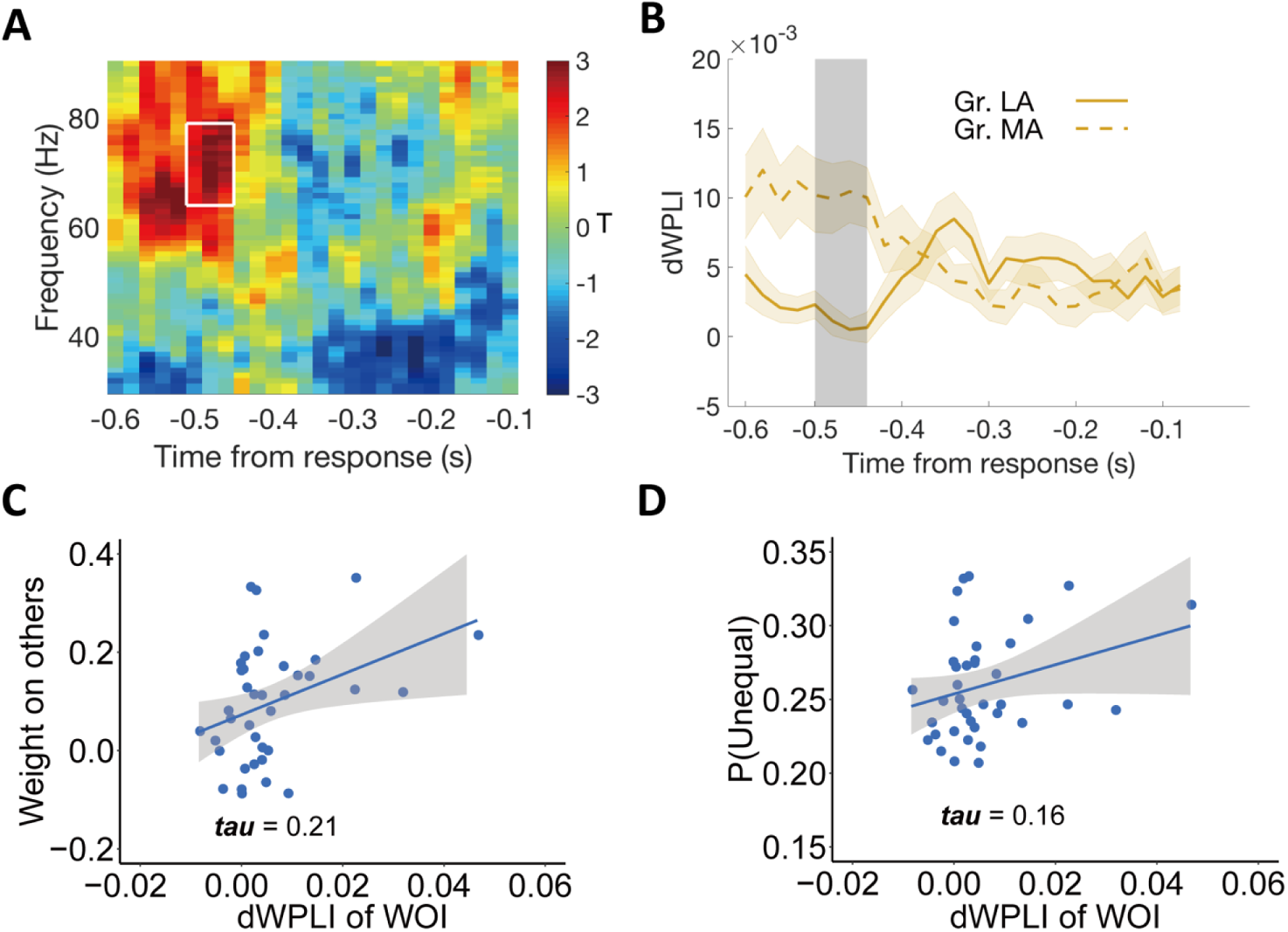
Individual differences in altruism relate to frontal-parietal synchronization. (A) Heatmap showing T-statistics for the differences between the more altruistic (MA) minus less altruistic (LA) group in phase coupling (dWPLI) between the frontal cluster shown in Figure 5A and the parietal cluster shown in Figure 2C, in the DIS context. A significant effect was identified in the gamma-band frequency range (~64 – 79 Hz) at the time window of ~ 520 to 460 ms before response (highlighted in white box). (B) Temporal dynamics of the average dWPLI strengths in the ~64 – 79 Hz frequency range. Grey shaded area indicates the duration of the significant effect. Colored shaded areas indicate ± 1 SEM. Gr., group. Significant clusters surviving the cluster correction for multiple comparisons at *p* < 0.05 are reported. (C) Correlation between the strength of frontal-parietal synchronization (dWPLI) in the identified time-frequency cluster and the OU parameter of weight on others. (D) Correlation between the strength of frontal-parietal synchronization (dWPLI) in the identified time-frequency cluster and the probability to choose unequal option in DIS context.

To visualize the hypothesized relationship between this neural effect and both model-based altruistic preferences and model-free unequal choice, we extracted the individual effect size in the DIS context from this temporal-frequency cluster and correlated it across participants with the weight on others’ payoffs (ω) and with the individual probability to choose the unequal option (note again that these analyses are for illustrative purposes and do not constitute circular inference). As expected, the strength of gamma-band synchronization was clearly related to the weight on the others’ payoff (*Kendall’s tau* (37) = 0.21, Figure 6C) and to the frequency of choosing the unequal option in the disadvantageous context (*Kendall’s tau* (37) = 0.16, Figure 6D).

The corresponding analyses for the advantageous context did not reveal any significant temporal-frequency cluster (see Fig. S13), which is in line with the findings that in this context, there was also no relation between individuals’ altruistic preferences (i.e., weight on others’ payoff) and ERP responses (see above). Taken together, these results suggest that frontal-parietal gamma synchrony may serve to incorporate information about other-payoff in evidence accumulation processes to promote altruistic preferences under disadvantageous inequality. Moreover, they suggest that while distribution decisions in different contexts and individuals are governed by comparable neural decision processes, individual differences in altruism may not only originate from differences in the perception processes but also from how this payoff-related information is shared between different regions by neural coherence.

## Discussion

The psychological and neural mechanisms underlying altruism are debated intensely. On the one hand, evolutionary theories suggest that relatively simple, unified mechanisms may have evolved to facilitate actions that run counter to our self-interest to benefit others (Gintis et al., 2003; Piliavin and Charng, 1990). On the other hand, models from economics and psychology have suggested that altruism is complex and may draw on a whole range of different motives that can be differentially triggered in different contexts and different individuals (Bester and Güth, 1998; Côté et al., 2015; Hein et al., 2016). Here we investigated - using computational modelling and EEG methods with high-temporal resolution – what cognitive processes that temporally evolve during the decision drive contextual and individual differences in altruism. This allowed us to investigate whether these differences reflect use of fundamentally different neural decision processes, or biases in the initial processing in the choice-relevant information before this information is integrated into a common decision process.

Our findings are more compatible with the latter: We find that a comparable response-locked parietal evidence accumulation process is deployed across different contexts and in different individuals, but also considerable differences in how the choice-relevant information is initially processed. Context differences were most evident in neural inequality signals related to self-payoff processing, whereas differences between individuals were most apparent in varying neural processing of other-payoffs. Thus, contextual and individual differences in altruism do not reflect use of fundamentally different neurocognitive choice mechanisms, but rather differences in how choice-relevant information is initially perceived and/or attended to.

### Neural sources of variability in altruism

Our study provides critical insights into the nature of altruism. Since altruism can co-occur with different types of motivations under different circumstances (Hein et al., 2016; Vekaria et al., 2017), it is crucial for us to pin down whether these motivations are expressed as fundamental differences in how choices are taken, or as variations in initial (perceptual/attentional) information processing. Either of these possibilities would have different implications for our understanding of the nature of human altruism, why people differ in their altruistic tendencies, and how these tendencies may be altered by interventions (Gao et al., 2018; Güroğlu et al., 2014; Jiang et al., 2016; Krajbich, 2018; Teoh et al., 2020; Tricomi et al., 2010).

Previous neuroimaging studies have been ambiguous on this point, showing that both common and distinct brain networks are active during altruistic decision making in different inequality contexts (Gao et al., 2018; Morishima et al., 2012; Yu et al., 2014). However, it is largely unclear what specific neuro-cognitive mechanisms underlie these different patterns of brain activation. Our findings fill this gap, by revealing that altruistic choices in different contexts are guided by a comparable neural decision mechanism that is well described by sequential sampling models; this mechanism is very similar to neural mechanisms reported to be recruited for perceptual and non-social value-based decision making (Pisauro et al., 2017; Polanía et al., 2014). Thus, our results support the view that altruism is not a functionally distinct aspect of human behavior but rather controlled by processes that are tightly integrated with general control processes required for non-social aspects of our actions (Krajbich et al., 2015).

Contextual and individual differences in altruism appear much more related to perceptual processing of the choice-relevant information, and thus how this information is differentially perceived and/or attended to before being integrated into the single brain mechanism. It has been long established in the decision making literature that attention to specific decision-relevant information can determine individuals’ final choices (Ghaffari and Fiedler, 2018; Krajbich, 2018; Krajbich et al., 2010) and even shape social preferences (Jiang et al., 2016; Teoh et al., 2020). For example, higher time pressure will bias gaze towards stimuli relevant for ones’ own interests, and the amount of this bias can predict changes in altruistic preferences across individuals (Teoh et al., 2020). Our neural results suggest perceptual or attentional origins for contextual and individual differences in altruism even under normal situations without time pressure. These differences relate not just to influences on the perceptual processing of self- or other-payoffs (Hutcherson et al., 2015) but also to the neural processing of inequality in different contexts (Yu et al., 2014). This extends previous reports that attention can affect processing of self-interest (Teoh et al., 2020) to neural processing of others’ interests and to how this information is communicated between different brain regions.

### Parietal evidence accumulation extends to social decisions

Our findings also enhance our understanding of the generality of parietal evidence accumulation signals for different aspects of human decision making. The involvement of parietal regions in evidence accumulation processes was initially established by perceptual decision making studies with electrophysiology and EEG signals (e.g., direction discrimination task), in which the build-up rate of neural activity in parietal regions was considered as a model-free measure of EA process (Brosnan et al., 2020; Churchland et al., 2008; Kelly and O’Connell, 2013; Kiani and Shadlen, 2009; O’Connell et al., 2012; Shadlen and Newsome, 2001). Recently, a similar role of parietal regions in evidence accumulation during value-based decision making was proposed by EEG studies of food choices based on subjective preferences (Pisauro et al., 2017; Polanía et al., 2014). One may wonder whether these neural signals may reflect low-level motor planning, rather than high-level decision mechanism linking decision values to motor responses. However, we believe this is not plausible, for two reasons. First, a recent simultaneous EEG-fMRI study has identified that the parietal evidence accumulation EEG signals can be traced to activity in posterior medial frontal cortex, which functions with value processing regions (i.e., ventromedial prefrontal cortex and striatum) to link decision values to motor responses, rather than reflect pure motor preparation (Pisauro et al., 2017). Second, since basic motor planning signals are typically observed in the hemisphere contralateral to the response hand (Kornhuber and Deecke, 2016; Schurger et al., 2021), the action readiness potential should be distributed more widely at left hemisphere (all participants responded with the right hand). Such signals related to motor planning should thus exhibit fundamentally different spatial-temporal characteristics than the parietal evidence accumulation signals observed here. Moreover, the causal role of parietal regions in evidence accumulation underlying decision making has been established in studies with both animals and humans (Licata et al., 2017; Polanía et al., 2015; Yao et al., 2020; Zhong et al., 2019; Zhou and J., 2019). Inactivation of projections from parietal regions to sensory specific regions (e.g., auditory cortex) will not only prolong rats’ evidence accumulation processes but also impair their decision performance (Yao et al., 2020; Zhong et al., 2019). Consistent with this, disrupting synchronization between parietal and frontal regions was also found to causally decrease the precision of value-based decision (Polanía et al., 2015).

In the current study, we not only revealed comparable evidence accumulation signals in parietal region in both inequality contexts, but also validated the tight relation of the model-free ERP build-up rate with the SSM model-based parameter (i.e., drift rate) for altruistic decision making. This correlation relates the parietal EEG signals to the decision mechanism captured by the SSM, and also confirms that the observed response-locked EEG signals are involved in linking decision values to responses, rather than just in pure motor planning (Kelly et al., 2021; Pisauro et al., 2017; Polanía et al., 2014; Schurger et al., 2021). Since both SSM model-based neural EA signals and model-free CPP signals reflect relatively late neural processes that are closer to the ultimate responses than the initial perceptual processing, our results do not exclude the possibility that different systems are involved in perceptual processing of the choice-relevant information at earlier stages. In fact, our findings provide evidence that although there may be comparable final decision processes guiding behavior in different contexts, these processes may receive inputs from different earlier neural processing of stimulus, and that these mechanisms may differ in different contexts. Thus, it appears that even high-level social decision making involves parietal evidence accumulation signals, rendering these signals a close approximation of a unified decision mechanism that can flexibly draw on many different types of information to guide choice (Kumano et al., 2016). Future studies using brain stimulation techniques may be needed to establish the causal role of parietal regions in evidence accumulation of social decision making.

### Inequality processing is crucial for altruistic decisions in disadvantageous and advantageous contexts

Although it is often observed that attention to specific aspects of decision-relevant information can bias individuals’ decisions (Jiang et al., 2016; Krajbich et al., 2010; Maier et al., 2020; Teoh et al., 2020), it is still largely unknown whether these biases reflect enhanced behavioral impact of single dimensions of choice-relevant information (i.e., here self-payoff or other-payoff), or of higher-level integration of various stimulus aspects (i.e., here inequality as the difference in self- and other-payoff). Both these possibilities have been incorporated in computational models of the evidence accumulation process, with little decisive evidence to back up either option.

Consistent with behavioral and modelling results, our EEG results suggest that integrated inequality processing, rather than simple perceptions of either self-payoffs or other-payoffs, plays a fundamental role for altruistic decision making in both inequality contexts. The inequality of payoffs between the two parties is detected early after stimulus onset (~ 240 – 360 ms), the conflict between inequality level and self-/other-payoff change is represented again at a late time window (~ 440 – 800 ms after stimulus onset). Both these effect show different temporal dynamics in different contexts, and they occur at similar time windows as typical ERP components related to anticipation, cognitive control, and attentional processing (e.g., N2 and FRN lasting ~ 200 – 300 ms, P300 peaking at ~ 300 – 400 ms, or P600 lasting occurring ~ 400 – 800 ms post-stimulus onset (Coulson et al., 1998; Sutton et al., 1965)). Our finding that inequality affects neural processing during both these time-windows argues that inequality is a fundamental motivational component that is computed by the brain already during early stages of perception, and is affected by subsequent attentional processing to bias choice.

Recent value-based decision making research also showed a late positivity response in a similar time window, varying with the value of stimulus attributes relevant to decision making (Harris et al., 2018). These studies consistently suggested that a stronger late positivity is evoked when stronger attention or cognitive resources are required to process stimuli with higher complexity or saliency. In the current study, the effects in the late positivity responses for inequality processing imply that this component does not necessarily encode the value of self-payoff, but participates in the processing of inequality, with different temporal dynamics across the two contexts. This temporal difference in neural signals may also account for the observation of shorter reactions times in the ADV than in DIS context, with faster conflict resolution associated with shorter response times (Chen and Fischbacher, 2020). Another potential (but more speculative) explanation for these findings could be people may differentially re-orient attention to different option attributes. That is, people may generally attend first to their own payoffs, but more altruistic people may then be more likely to attend others’ payoffs (Teoh et al., 2020), so that participants may be differentially taxed in different contexts by the “meta-decision” of where to direct attention to guide their decisions. Although we ruled out eye moverments in the current study, participants in the advantageous context may process their own payoffs faster and re-orient attention to others’ payoffs earlier than in the disadvantageous context, where they may need to take more time to process their (lower) own payoffs. Since we did not manipulate or measure attention orientation in the current study, we cannot arbitrate between these different explanations for the observed effects, which may perhaps be further investigated by eye-tracking studies. Irrespective of these possible explanations, our findings again underline that the processes involved in controlling altruism in different inequality contexts differ mainly in terms of how the decision-relevant information guiding choice is perceived and attended to (Gao et al., 2018; Güroğlu et al., 2014; Yu et al., 2014).

### Individual differences are linked to information processing

A core question in research of altruism has always been why people differ in their altruistic motivation and behavior (Gao et al., 2018; Morishima et al., 2012; Yu et al., 2014). Our EEG results add a novel perspective to this literature, by showing that individual differences in altruistic preferences (i.e., the decision weight placed on others’ payoffs) is related to early brain processing and fronto-parietal information exchange of the information quantifying other-interest. While this highlights that perceptual and attentional processing of choice-relevant information may strongly influence choice outcomes even for high-level social choices, it is noteworthy that individual differences in altruism were linked to processing of different aspects of the choice situation (others’ outcomes) than those aspects that seemed related to contextual differences (inequality in outcomes). This suggests that different perceptual/attentional biases underlie variations in altruism across situations and people, even though these variations may at first glance appear to be of similar magnitude. Thus, our results demonstrate more generally that neural measures can help us to better understand the fundamental motivations underlying variability in behavior across people and contexts, which would be impossible to achieve with behavioral measures alone (Hein et al., 2016).

Our finding that fronto-parietal gamma-band coherence correlated with altruism identifies a specific neural mechanism that may underlie how the information of other-interest is integrated into choices. This process shows clear parallels with previous findings from perceptual and non-social value-based choices, where related increases in gamma-frequency band coherence have also been shown to facilitate information transfer between distant brain regions to improve performance (i.e., precision) (Gregoriou et al., 2009; Polanía et al., 2014; Vinck et al., 2013; Womelsdorf et al., 2006). Specifically, frontal regions involved in assigning value to choice options have been found functionally couple with parietal regions in the gamma band to implement evidence accumulation to value-based social decision making (Basten et al., 2010; Philiastides et al., 2010; Pisauro et al., 2017), and one study even demonstrated that stronger frontal-parietal phase coupling in gamma band can causally increase precision of value-based decision making (Polanía et al., 2015). Based on these earlier findings and our present results, it appears possible that stronger information transfer or sharing of others’ payoff between frontal and parietal regions can indeed account for greater altruistic preferences in social decision making. Nevertheless, such modulation of frontal-parietal interregional information transfer on altruistic preferences may be relayed by subcortical regions, as suggested by a recent monkey study showing that stronger gamma-band synchronization between anterior cingulate cortex (ACC) and basolateral amygdala is related to enhanced other-regarding preferences (Dal Monte et al., 2020). Given the limitation of spatial resolution in EEG recordings, we cannot pinpoint the involvement of specific cortical or subcortical regions, but regions like ACC and amygdala could be candidate regions involved in information transfer underlying altruistic preferences in humans as well.

It is noteworthy that we only observed modulation of local and large-scale processing of other-payoff change in the DIS context, but not in the ADV context. One potential explanation for this context differences is that individuals’ relatively disadvantageous position in the DIS context may make selfish people more inclined to upward social comparison, and thus to higher sensitivity for profit change of others who are already better off than themselves (Boyce et al., 2010; Payne et al., 2017). The ADV context may more uniformly bias participants to put a stronger weight on others’ profit - as suggested by our modelling results and previous studies (Morishima et al., 2012) - thereby potentially reducing individual difference in sensitivity to others’ profit change.

Our study also suggests that the effects of inequality contexts on the decision process may relate to framing effects that are tightly related to how information is presented to participants (Dietze and Craig, 2021). In the current paradigm, the two allocation options were presented sequentially, so that participants knew the context they were in before the EA process started with the presentation of the second option. This may explain why the inequality contexts affected altruistic decisions not just by modulating perceptual processing, but also by changing other latent decision processes such as criterion required for a decision to be taken (i.e., decision threshold). The higher payoffs in ADV context may put participants in a relative gain frame and render them more impulsive (i.e., lower decision threshold), since they benefit more than the others from both options, whereas the lower payoffs in the DIS context may put participants in a relative loss frame and drive them to be more cautious (i.e., higher decision threshold) and integrate more evidence to make the final decision (Diederich and Busemeyer, 2006; Dietze and Craig, 2021). Importantly, our experiment setup is similar to situations in real life, since people usually know whether they have more or less money than others when deciding whether to behave altruistically. Nevertheless, our results emphasize that future studies should examine how different information presentation formats may bias different latent decision processes, such as attention to different aspects of information (e.g., self-payoff, other-payoff, or inequality level) and impulsivity, to influence social preferences and decisions, which has rarely been investigated in previous studies (Gao et al., 2018; Hutcherson et al., 2015; Morishima et al., 2012).

### Facilitation of altruistic behavior

Our findings provide empirical evidence for improving strategies to promote altruism from the perspective of decision processes. Most previous studies focused on clarifying the relationship between individual variations in altruistic preferences and dispositional empathy (i.e., empathy-altruism hypothesis) and suggested that facilitating individuals’ empathic concerns for others will enhance altruistic preferences or behavior (Batson et al., 2007; FeldmanHall et al., 2015; Hein et al., 2010; Lockwood et al., 2017). However, it is possible that corresponding interventions meant to enhance empathy do not operate mainly by enhancing emotional concern for others but rather by shifting attention to corresponding information. This would fit our findings that variations in altruism across different contexts and individuals may have more perceptual or attentional origins, and suggests that it may be possible to have substantial influences on altruistic behavior by interventions that shift attention to related choice-relevant information even without explicitly cuing or training empathic concern (Jiang et al., 2016; Teoh et al., 2020).

Our approach to examine altruism across contexts can also help to improve diagnosis and treatment for socially apathetic people or other people with high subclinical or clinical level of psychiatric and neurological disorder (e.g. psychopathy, autism, or alexithymia), by investigating which latent components of decision processes underlying altruistic behaviors may be altered in these disorders and changed by corresponding treatments (Bird and Viding, 2014; FeldmanHall et al., 2013).

Taken together, the current study provides a unified neural account of contextual and individual differences in altruism, by demonstrating that comparable neural evidence integration process underlies variation in altruistic decisions across different inequality contexts and individuals (which is often captured by different parameters in econometric models). These decision processes recruit both local neural activity in parietal cortex and large-scale neural synchronization between different brain regions. Variations in behavior did not relate to fundamental differences in these neural decision mechanisms, but rather to how the choice-relevant information feeding into them is processed during earlier stages of perception. Thus, the SSM-EEG framework we employed here provides a powerful tool to clarify whether changes in altruistic preferences across contexts and individuals reflect fundamentally different motives and choice processes, or rather more perceptual/attentional effects that may be amenable by corresponding interventions. More generally, our study suggests that this approach may be employed to study neural origins of individual and contextual variations also in other types of social decision making that need to integrate value signals from multiple sources, such as empathy-based or reciprocity-based altruism, moral decisions, and sophisticated strategic decision making.

## Materials and Methods

### Participants

Forty-one participants (16 females) participated in the study. Participants were informed about all aspects of the experiment and gave written informed consent. All the participants were right-handed, had normal or corrected-to-normal vision, were free of neurological or psychological disorders, and did not take medication during the time the experiment was conducted. They received between 75 – 122CHF (depending on the realized choices) for their participation. Three participants were excluded from analyses due to excessive EEG artifacts (remaining 38 participants: 19 – 31 years of age, mean 24.5 years). Sample size was determined based on previous study (Hutcherson et al., 2015) that detected a significant DDM parameter of weight on others with an effect size of d = 0.4. We thus determined our sample size based on d = 0.4 with G*Power 3.1, which suggested that we need 41 participants to have adequate power (1 – β > 0.80) to detect an effect with d = 0.4 at the level of α = 0.05. The experiment conformed to the Declaration of Helsinki, and the protocol was approved by the Ethics Committee of the Canton of Zurich.

### Task

Each participant made 416 real decisions in a modified Dictator Game, requiring them to choose one of two allocations of monetary amounts between the participant and a receiver (another participant in the same study). We used two inequality contexts: In the disadvantageous context (DIS), the token amounts for participants were always lower than for the receiver; in the advantageous context (ADV), they were always higher. On each trial, one of the two options was revealed at the beginning of each trial (i.e., the reference option) and the other option was revealed at a later time (i.e., the alternative option, see below and Figure 1A for details). There were 4 levels of reference options (i.e., 24/74, 24/98, 46/74, and 46/98) in each context. For the DIS context, the numbers before the slash denote the token amounts allocated to the participant and the numbers after the slash the token amount given to the partner, and vice versa for the ADV context. There were 26 alternative option levels corresponding to each reference option level, with half of them being more equal than the reference option and the other half being more unequal than the reference option. The token differences for each party between the alternative and the reference options ranged from −19 to 19. To make sure that our trial set is able to capture a large range of altruistic preferences and can properly estimate the parameters of the Charness-Rabin model, we not only included decisions requiring opposite changes in self- and other-payoff (i.e., maximizing one and minimizing the other), but also a small number of choice options leading to changes in inequality but increases or decreases of both self- and other-payoffs. To avoid repetitions of exactly the same choices, we included two different trial sets with the same reference options and similar distributions of alternative options. The second trial set was generated by adding a random jitter (i.e., −1, +0, or +1) to self-/other-payoffs of the alternative options of the first trial set (Fig. S1).

At the beginning of each trial, participants saw a central fixation cross together with a reference allocation option on one side of the cross. The alternative option, to be shown on the opposite side of the fixation cross, was initially hidden and replaced with “XX” symbols. Participants were asked to keep their eyes fixated on the central cross for at least 1 s (this was controlled by the use of eye tracking, see below). Only after successful fixation for at least 1 s, the font color for all the stimuli changed from blue to green (the change direction of font color was counterbalanced across participants), indicating the initiation of the trial. After a temporal jitter of 0.5 – 2 s (uniform distribution with a mean of 1.25 s), the alternative option was revealed (with XX symbols replaced with actual amounts). Participants had to choose the left or right option by pressing the corresponding keys on the response box with right index or middle finger within 3 s. The selected option was highlighted with color change. Note that since the fixation was constrained during the whole trial before participants’ response (for more detailed description, see the section on Eye tracking), the retinal processing of the first and the second option should be the same. Since all payoff stimuli were presented close to the fixation cross (i.e., visual angle smaller than 3°), participants were able to see the numbers clearly without shifting their gaze.

The task was divided into 4 blocks, with each block lasting around 10 minutes. In total, the experiment took around 50 minutes to 1 hour, including breaks between blocks.

To avoid potential experimental demand effects, we did not explicitly describe the classification of decision contexts as advantageous or disadvantageous in the instructions. Nevertheless, since participants were presented with the first option at the beginning of each trial, they were clearly aware of whether they were better or worse off than the other at the beginning of each trial. Such a sequential presentation resembles real life contexts in which people are usually already aware of whether they are in better (advantageous) or worse (disadvantageous) position, before having to decide whether to improve the welfare of the other.

To avoid potential attentional or visual processing bias due to fixed position of reference/alternative payoffs, we counterbalanced stimulus positions (left versus right) trial-by-trial within participant. To lower processing load and avoid potential response errors due to mis-recognition, we fixed the position of self/other payoffs within participant and counterbalanced it between participants.

The participation payment was determined at the end of the experiment and consisted of three parts: a base payment of 45 Swiss Francs (CHF, around $45 at the time of the experiment), one bonus (Bonus A) payment that depended on participants’ own decisions, and another bonus (Bonus B) payment that was determined based on the choices of previous participants whose outcomes were allocated to the current participant. To determine these bonus payments, the participant drew two envelopes from two piles of envelopes, one pile for bonus A and one pile for bonus B. Each envelope contained five different randomly determined trial numbers.

For Bonus A, the numbers in the envelope were the trial numbers that would be selected from the full list of the participant’s choices to be paid out. We calculated the mean payoff from these options and paid this as the first bonus.

For Bonus B, the numbers in the envelope were chosen from the full list of choices taken by previous participants. This list of choices was randomly drawn from the full list of choices of all previous participants, so that the participant was randomly paired with a different person on every round. The mean of the partner’s payoffs across the chosen five rounds was paid out as the second bonus.

The partner’s payoffs resulting from the participant’s own choices also entered the full list of choices for future participants, meaning that they were be paid out to these participants if any of the current participant’s choice was selected. The exchange rate was 1 token = 0.5 CHF.

### EEG recordings

EEG signals were recorded from 128 scalp sites using sintered Ag/AgCl electrodes mounted with equidistant hexagonal layout using a Waveguard Duke 128 channels cap (http://www.ant-neuro.com/). The cap was connected to a 128-channel QuickAmp system (Brain Products, Munich, Germany). All EEGs were referenced online to an average reference electrode. Electrode impedance was kept below 5 kΩ for all the electrodes throughout the experiment. The bio-signals were amplified with a band pass from 0.016 to 100 Hz and digitized on-line with a sampling frequency of 500 Hz. The EEG cap was set up on each participant’s head before he/she entered the soundproof and electromagnetically shielded chamber to perform the decision-making task during EEG recordings.

### Eye tracking

We used an EyeLink-1000 (http://www.sr-research.com/) to track and record participants’ fixation patterns with a sampling frequency of 500 Hz. Before each trial, participants were instructed not to blink and keep their eyes fixated at the central fixation cross for 1 s before the trial started. From the start of the trial until the response was detected, participants were also asked not to blink and maintain fixation (tolerance 3°). If they failed to do so, the trial was aborted, and participants were informed with a message reminding them that the trial was invalid due to eye movements. Such trials were discarded in the EEG analyses. Participants practiced the task and fixation in a 10 min practice session. For this practice session, participants were presented with a different set of allocation options than those used in the real experimental session.

### Computational model

We used the Ornstein-Uhlenbeck (OU) process to model evidence accumulation underlying individuals’ decisions. The OU process updates the evidence (EA) at each subsequent time step *t* with the following equation:

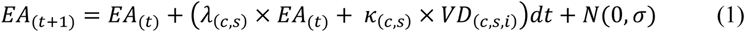

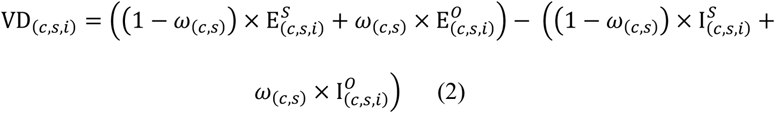

with indices c for conditions (c = DIS for disadvantageous inequality context, c = ADV for advantageous inequality context), s for participants (s = 1, …, N_participants_), and i for trials (i = 1, …, N_trials_). 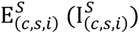 indicates participants’ payoff of the equal (unequal) option in condition c, for participant s and trial i; 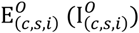 indicates the partners’ payoff in the equal (unequal) option in in condition c, for participant s and trial i.

The following free parameters were estimated for each participant and context separately:

*α*_(*c,s*)_ : decision threshold, which indicates the amount of evidence required for making a decision (symmetric around zero);

*β*_(*c,s*)_: starting point, which represents the initial bias towards equal or unequal option (0 = no bias);

*κ*_(*c,s*)_: drift rate modulator, which scales the input of the subjective value difference between the equal and the unequal option (VD);

*ω*_(*c,s*)_: relative weight on others’ payoffs, which reflects the individual’s concern for others’ profits (i.e., altruistic preferences);

*λ*_(*c,s*)_: leak strength, which adaptively controls acceleration or deceleration of evidence accumulation at the current time point given the amount of evidence at the previous time point (Bogacz et al., 2006; Brunton et al., 2013);

*τ*_(*c,s*)_: non-decision time (nDT), which accounts for sensory and motor processes not contributing to evidence accumulation per se, such as early visual processing and motor-response initiation.

Evidence accumulation starts with an initial value *EA*_(0)_ equal to the starting point parameter (*β*). This value is then updated in discrete time steps of *dt* = 0.001 s until |*EA*_(*t*)_| is greater than the decision threshold parameter (*α*). The model will make an equal decision when *EA*_(*t*)_ > *α*, and make an unequal decision when *EA*_(*t*)_ < −*α*. The noise of evidence accumulation at each time step is drawn from a Gaussian distribution of *N*(0, σ), and we fixed σ as 1.4. The subjective value differences between the equal option and the unequal option (VD) is constructed based on the Charness-Rabin utility model (Charness and Rabin, 2002), where 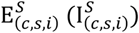 indicates participants’ payoff of the equal (unequal) option in condition c, participant s, and trial i, and 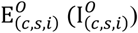 indicates the partners’ payoff of the equal (unequal) option in condition c, participant s, and trial i. Once the evidence (*EA*_(*t*)_) reaches the decision threshold, the RT is calculated as *t* × *dt* + *τ*, where *τ* is the non-decision time.

We used the differential evolution algorithm introduced by Mullen et al. (Mullen et al., 2011) to estimate the values of the free parameters described above separately for each participant and context. We ran the estimation with 60 population members over 150 iterations. For each iteration, we simulated 3,000 decisions and RTs for VD in each trial for each participant with the free parameters for each population member. The likelihood of the observed data was computed given the distribution generated by the 3,000 simulations for a certain set of parameters. Then the population evolves to a set of parameters maximizing the likelihood of the observed data with the procedures described by Mullen et al. (Mullen et al., 2011). We checked the evolution of the population across the 150 iterations and found that the algorithm reached a set of best-fitting parameters well before 150 iterations in our data. The upper and lower bounds of each parameter are listed in Table S4. We selected uninformative bounds of these parameters based on previous studies (Maier et al., 2020; Polanía et al., 2014, 2015). The bounds for the parameter of weight on others (*ω*) were set from −1 to 1 because these are the limits inherent in the model (the sum of the weighting on others’ payoff and ones’ own payoff is 1 in the Charness-Rabin model).

In the above OU model (full), we estimated all the parameters separately across contexts and participants. To confirm that this full OU model is the best model to explain individuals’ behaviors, we fitted a series of alternative models for model comparison analyses. Specifically, first, we fit a drift diffusion model (DDM) that kept the same parameters as the full OU model except the leak strength parameter. Second, to confirm that the weight on others (*ω*) and decision threshold (*α*) were indeed different across contexts, we included models in which the weight on others (*ω*) and/or decision threshold (*α*) were constrained to a unique value across contexts as benchmark models. Model comparison results showed that the full OU model was a better fit than the other models to account for participants’ behavioral data (i.e. with the lowest BIC = 4058 ± 46 (Mean ± SE)). For details, see Table S5. The full OU model was the best fit for all the 38 participants (100%). These results demonstrated that separating the parameters for leak strength, weight on others, and decision threshold across contexts did improve model fits while taking into account model complexity. In addition, we also estimated one OU model which used the 1^st^ vs 2^nd^ choice (OU order model), rather than the equal vs unequal choice, as the decision boundaries. This model also correctly captured response speed and choice (Fig. S14), and the estimated parameters (i.e., weight on others, drift rate, decision threshold, and non-decision time) were highly correlated between the OU order model and the winning model in both contexts (Fig. S15). The BIC of the OU order model (4059 ± 47) was slightly higher than the winning model. Since we are interested in the decision mechanism of individuals’ altruistic choice, rather than the effect of offer presentation order, the OU order model was only considered as a validation of the winning model.

To evaluate model performance, we also calculated the sensitivity/specificity and balanced accuracy of the OU model. These measures are defined as follows:

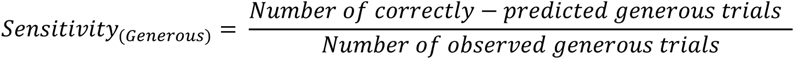

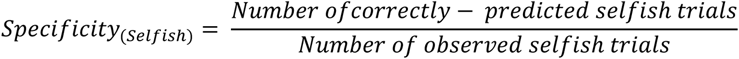

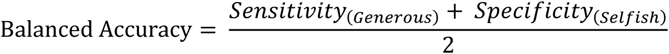

Please note that while the model exhibited lower and more variable predictive accuracy for unequal (equal) choices in the DIS (ADV) contexts, this can be explained by how the different choices tap differentially into the individuals’ variable preferences: In each context, one choice option is dominant, as evident by that finding that P(equal) = 0.74 ± 0.006 in DIS and P(equal) = 0.27 ± 0.009 in ADV. Participants thus more uniformly choose the dominant equal (unequal) choice option in the DIS (ADV) context and only people with stronger altruistic preferences choose the non-dominant unequal (equal) choice option, leading to larger variability. However, this asymmetry cannot have biased our results, since we only included a single weighting parameter to capture individuals’ decision weight on other-payoff (i.e., altruistic preferences) in the model for each context, rather than having two different parameters to separately capture individuals’ preferences for equality and inequality. It is also important to note that there is no difference in overall predictive accuracy between ADV and DIS conditions.

### EEG analysis

We performed EEG data analyses using Fieldtrip (Oostenveld et al., 2011) implemented in Matlab R2016b (MathWorks). We first identified eye movements and other noise artifacts using independent component analysis, and removed artifact components based on careful inspection of both topography and power spectrum of the components. We then used discrete Fourier transform to filter line noise. For response-locked analyses, epochs were extracted for a 1,200 ms time window around the response, ranging from 1,000 ms before to 200 after the response timepoint. For stimulus-locked analyses, we extracted epochs for a 1,200 ms time window around the 2^nd^ option onset, ranging from 200 ms before to 1,000 ms after the stimuli onsets. Next, we visually inspected each individual trial and removed all trials with extremely high variance (e.g., muscle artifacts) from the data. We excluded three participants from the analysis due to excessive artifacts. For the remaining 38 participants, 21% ± 8.5% (Mean ± SD) of the trials were rejected.

### Response-locked EEG analysis

To validate evidence accumulation (EA) processes with EEG data, we used an analytical approach previously established in our lab (Polanía et al., 2014). We first simulated EA curves for each trial 500 times using each participants’ best-fitting parameters and self-/other-payoffs in each trial and averaged these 500 simulated curves to compute a trial-specific model predicted EA trace.

For EEG data, we focused on a time window starting 600 ms prior to the response and lasting until 100 ms before the response. We excluded the last 100 ms before the response to avoid motor execution-related signals due to abrupt increase in cortico-spinal excitability during this period. We examined the EEG-signal-associated EA processes using a cross-validation approach which combined both qualitative and quantitative criteria.

We divided trials into even and odd trials to test predictions of the EA model for each participant. We first used the even-numbered trials to identify channels where EEG signals were closely related to the shape of the model-predicted EA signal in each context, and then formally tested the model predictions against the data from the independent odd-numbered trials.

We carried out the analysis for the even-numbered trials in two steps: First, we calculated a linear regression with 250 time points (i.e., 600 ms to 100 ms in steps of 2 ms before response in each trial) × N trials (i.e., number of trials in each context) between the ERP magnitude of even-numbered trials and model-predicted EA signals. Channels that survived Bonferroni correction at *p* < 0.05 were selected. Then, from the channels surviving the quantitative test, we further selected channels for which the average magnitude laid within 1 SE of the model predictions to ensure that the ERP signals were closely related to the EA model predictions across the full temporal interval.

Then, to test cross-validation predictions, we examined the relationship between the model-predicted ERP signals and actual ERP signals in the fully independent odd-numbered trials, based on the channels identified in the previous analysis.

To compare the relationship of model predictions and data between DIS and ADV contexts, we performed a *t* test of the difference between contexts in correlation (*r_Pearsons_*) between model predictions and data for each EEG channel.

To account for multiple comparisons, we identified significant clusters based on two-sided *t* statistics with a threshold of *p* < 0.05 at channel level and at least 3 significant neighbouring channels in a single cluster. For each cluster that survived the threshold, the size of the cluster was defined as the integral of the t scores across all the channels of the cluster and its significance was tested with a permutation statistic (i.e., performing iterations 5,000 times with shuffled context labels to generate a distribution of cluster sizes for the null hypothesis of no difference between contexts) (Maris, 2012). Significant clusters surviving the cluster correction for multiple comparisons at *p* < 0.05 are reported. We applied this threshold for all the whole-brain EEG analyses to identify significant clusters in the current study.

To calculate the built-up rate, we followed Kelly and O’Connell’s work (Kelly and O’Connell, 2013) and extracted grand average response-locked ERP waveforms from the identified clusters in previous analysis. Then, we fitted a linear slope to the waveforms in the time window of −300 ms – −100ms before response and took the steepness of these slopes as indices of build-up rates for each participant.

### Stimulus-locked EEG analyses

With stimuli-locked EEG analyses, we aimed to identify channels and time windows in which neural activity is associated with self- and other-interest (i.e., self-/other-payoff change between the 2^nd^ and the 1^st^ option) processing in each context. For each channel, each epoch was integrated over 40 ms windows from 0 to 1000 ms after the 2^nd^ option onset, generating a matrix of EEG signals in 128 × 25 time windows for each trial. Then, we used the EEG signals across all the trials in each context in a linear regression model described as follows:

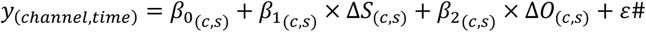

where *y*_(*channel*,*time*)_ is a matrix of trial-by-trial data for each channel and each time window, Δ*S* is the matrix of standardized participants’ own payoff changes from the 1^st^ to the 2^nd^ option (2^nd^ minus 1^st^) across all the trials, and Δ*O* is the matrix of standardized partners’ payoff changes from the 1^st^ to the 2^nd^ option across all the trials. This linear regression produced a set of estimated coefficients (i.e., *β*_1_ and *β*_2_ maps) measuring the parametric effect strengths of Δ*S* and Δ*O* on neural activity for each channel and time window in each context and participant (*c* for condition, and *s* for participant).

To investigate how the neural weighting of Δ*S* and Δ*O* varied across DIS and ADV contexts, we contrasted *β*_1_ (*β*_2_) maps between the two contexts by performing paired-sample *t* tests over all participants. We report the results based on permutation tests to account for multiple comparisons as described above. Significant clusters surviving the cluster correction for multiple comparisons at *p* < 0.05 are reported.

To further investigate whether and how the neural weighting of Δ*S* and Δ*O* varied across individuals with different altruistic preferences, we split participants into more altruistic (MA) and less altruistic (LA) groups by the median of the weight-on-others parameter or of the starting point, derived from the OU model in each context, respectively. For each context, we contrasted *β*_1_ (*β*_2_) maps between the two groups by performing independent-sample t tests. To visualize the effects of significant clusters and to investigate EEG signals associated with processing of different dimensions of information (i.e., Δ*S* and Δ*O*), we further split trials in each context into trials with positive and negative Δ*S* or trials with positive and negative Δ*O* and extracted average ERP waveforms for each of these sub-conditions. For the interaction analyses of payoff change and context over ERP waveforms, we also report the results based on permutation tests to account for multiple comparisons as described above. Only significant clusters surviving the cluster correction for multiple comparisons at *p* < 0.05 are reported.

### Connectivity analysis

To investigate the neural communication characteristics between channels associated with processing of other-payoff and channels associated with evidence accumulation processes, we performed connectivity analyses using the debiased weighted phase lag index (dWPLI) (Vinck et al., 2011). This coherence statistic is based only on the imaginary component of the cross-spectrum, which ensures that potential differences in power between the contexts or groups do not affect direct cross-context or cross-group comparisons of coherence. It also removes the type of bias due to small sample size in estimation of the phase lag index and is less sensitive to noise and volume conduction.

We first performed spectral estimates (both power and cross-spectral density) for each response-locked epoch using a multi-taper method implemented in Fieldtrip. The time-frequency analyses were performed in the frequency ranging between 16 and 100 Hz. The length of the temporal sliding window was eight cycles per time window in steps of 0.02 s. The width of frequency smoothing was set to 0.3×f with a frequency resolution in steps of 1 Hz. We then calculated the dWPLI of all possible pair-wise connections between the Δ*O* processing channels in DIS shown in Figure 5A and the EA-related channels in DIS shown in Figures 2C for each participant. We focused on the signals ranging between 600 ms and 100 ms before responses at gamma band frequency (30 Hz to 90 Hz).

To compare the dWPLI between MA and LA groups, we conducted an independent-sample t-statistic of dWPLI differences between groups for each channel and time step. We thresholded the t-statistic map with *p* < 0.05. For each cluster surviving this threshold, the size of each cluster was defined as the integral of the t scores (group effect) across all channels of the cluster and its significance was tested with a permutation statistic, i.e., the cluster identification was repeated 5,000 times by shuffling group labels to generate a distribution of cluster sizes for the null hypothesis of no difference between groups. Only significant clusters surviving the cluster correction for multiple comparisons at *p* < 0.05 are reported.

## Acknowledgements

This project has received funding from the European Research Council (ERC) under the European Union’s Horizon 2020 research and innovation programme (grant agreement No 725355, ERC consolidator grant BRAINCODES).

