## supplementary information for "A unified neural account of contextual and individual differences in altruism"

### SI Results

*Behavior: Choice depends differentially on self- and other-payoffs across contexts*

To visualize relationships between the effect of self-payoff, other-payoff, and inequality context on choice outcome identified in model-free linear mixed-effects regressions, we divided trials for each participant and context into seven bins based on the z-score of the Self-payoff Change (ΔS). Then we further split trials in each bin into trials in which the 2nd option would benefit or harm the other financially, relative to the 1st option (i.e., Increased other-payoff (OP) when the 2nd option profits the other versus Decreased other-payoff (OP) when the 2nd option leads to lower payoff for the other relative to the 1st option, Fig. S2A left and middle panels). We then fitted psychometric functions to these data and compared their slopes to determine how sensitive choices were to changes in self-payoff change in the two inequality contexts, and how this differed between choices that increased versus decreased the other person's payoff from 1st to 2nd option (Fig. S2A left and middle panels). For choices that would decrease the other's payoff, participants reacted less strongly to the associated possible increases in self-payoff in the advantageous context (lower slope of Self-payoff Change function in ADV than DIS for Decreased Other-payoff (OP) trials; 1.74 ± 0.12 vs 2.02 ± 0.10 (MEAN ± SE), ADV vs DIS: 95% CI [- 0.59 – 0.03], Cohen's d = - 0.41, $t(37) = -1.85$, $p = 0.07$). By contrast, for choices that would increase the other's payoff, people were equally sensitive to increases in self-payoff between both inequality contexts (Increased Other-payoff (OP); ADV: 2.09 ± 0.12, DIS: 2.03 ± 0.11, ADV vs DIS: 95% CI [-0.20 – 0.31], Cohen's d = 0.07, $t(37) = 0.45$, $p = 0.65$; direct comparison of both effect: $F(1,37) = 3.54$, $p = 0.068$, Fig. S2A right panel).

*Behavior: Presentation order of options does not affect equal/unequal choices*

To test whether properties of the reference option would affect individuals' choice, we ran a linear mixed-effects model in which individuals' equal/unequal choice (equal =1, unequal =0) was the dependent variable and the other-payoff difference between the equal/unequal option and the advantageous/disadvantageous context were independent predictors. Since the self-payoff difference between the equal and unequal option was negatively correlated with other-payoff difference, we did not include it in this control analysis. Importantly, we also included a predictor indicating whether the equal option was the reference (0) or the alternative option (1). This reference option indicator was not a significant predictor for individuals' equal choice (Table S2). Therefore, the asymmetry of the reference/alternative option or the presentation order of options did not affect participants' equal/unequal choice.

*ERP results*

To investigate how the brain processes payoff information, we analyzed ERPs related to different aspects of the payoff information (e.g., self-payoff/other-payoff changes between the 2$^{nd}$ option and the 1$^{st}$ option) upon stimulus presentation. In the analyses reported in the main text, we have identified two different spatial-temporal clusters showing opposite effects of self-payoff changes between different inequality contexts (i.e., earlier effect: from ~ 240 to 360 ms after stimulus onset, Figure 4A; later effect: from ~ 440 to 800 ms after stimulus onset, Figure 4C).

To further clarify whether the time courses for neural processing of self-payoffs were different or similar in the two contexts, we first directly compared the ERP components corresponding to the effects identified in the above analyses. For this analysis, we categorized trials in each context into trials in which the 2$^{nd}$ option increases self-payoff (i.e., trials with more equal 2$^{nd}$ option in DIS and more unequal 2$^{nd}$ option in ADV) and trials in which the 2$^{nd}$

option decreases self-payoff (i.e., trials with more unequal $2^{nd}$ option in DIS and more equal $2^{nd}$ option in ADV), and examined average ERP waveforms for each level of self-payoff change and context. Second, we performed interaction analyses between context (ADV vs. DIS) and self-payoff change (Increased SP vs. Decreased SP) over average ERP signals in the time window of ~440 to 800 ms after stimulus onset identified in the initial regression analysis.

In the following two sections, we reported detailed statistic information for these stimuli-locked ERP results.

*Different ERP responses are associated with self-interest in different contexts*

By directly comparing ERP dynamics corresponding to the effects identified in the regression analyses reported in the main text, we confirmed that for both time windows (i.e., the early window: ~ 240 to 360 ms after stimulus onset, and the late window: ~ 440 to 800 ms after stimulus onset), ERPs associated with the increase and decrease of self-payoff yielded opposite neural effects in the DIS and ADV contexts (see Figure 4B & 4D). Specifically, factorial analysis of ERPs revealed that for the earlier time window (~ 240 to 360 ms after stimulus onset), in the ADV context, self-payoff increases (-0.75 ± 0.15 μV, Mean magnitude ± SE, 95% CI [-1.04, -0.46], Cohen's d = -0.84, $t(37) = -5.17$, $p < 0.001$) evoked a stronger negative response than self-payoff decreases (-0.59 ± 0.13 μV, 95% CI [-0.86, -0.33], Cohen's d = -0.74, $t(37) = -4.59$, $p < 0.001$) at trend level (Increased SP vs. Decreased SP: 95% CI [-0.32, 0.01], Cohen's d = -0.31, $t(37) = -1.91$, $p = 0.06$). Similarly, in the DIS context, self-payoff decreases (-0.66 ± 0.15 μV, 95% CI [-0.95, -0.36], Cohen's d = -0.72, $t(37) = -4.46$, $p < 0.001$) evoked a more negative response than self-payoff increases (-0.54 ± 0.13 μV, 95% CI [-0.80, -0.29], Cohen's d = -0.70, $t(37) = -4.29$, $p < 0.001$) also at trend level (95% CI [-0.24, 0.02], Cohen's d = -0.28, $t(37) = -1.70$, $p = 0.098$, Figure 4B). In the later time window (~ 440 to 800 ms after stimulus onset), stronger positive responses were seen for self-payoff increases (1.05 ± 0.11 μV, 95% CI [0.83, 1.26], Cohen's d = 1.59, $t(37) = 9.81$, $p < 0.001$) than self-payoff decreases (0.86 ± 0.07 μV, 95% CI [0.72, 1.01], Cohen's d = 1.97, $t(37) = 12.17$, $p < 0.001$) in ADV (Increased SP vs. Decreased SP: 95% CI [0.04, 0.32], Cohen's d = 0.43, $t(37) = 2.64$, $p = 0.012$), whereas stronger positive responses were observed for self-payoff decreases (0.95 ± 0.09 μV, 95% CI [0.77, 1.13], Cohen's d = 1.75, $t(37) = 10.80$, $p < 0.001$) self-payoff increases (0.76 ± 0.08 μV, 95% CI [0.61, 0.92], Cohen's d = 1.61, $t(37) = 9.92$, $p < 0.001$) for DIS (Decreased SP vs. Increased SP: 95% CI [0.07, 0.30], Cohen's d = 0.53, $t(37) = 3.26$, $p = 0.002$, Figure 4D).

*Temporal difference of self-interest/inequality processing across contexts*

For the time window of ~ 440 to 800 ms after stimulus onset, we observed an earlier neural response to the self-payoff change in ADV (~ 450 to 600 ms after stimulus onset) than in DIS (~ 630 to 700 ms after stimulus onset, Figure 4D and 4E). To visualize the interactive effects of self-payoff change and context in these two windows of interest (WOIs), we extracted the ERP data from each WOI. Planned comparison showed that for the early WOI, self-payoff increases evoked stronger neural responses (1.04 ± 0.10 μV, 95% CI [0.84, 1.24], Cohen's d = 1.66, $t(37) = 10.26$, $p < 0.001$) than self-payoff decreases (0.71 ± 0.07 μV, 95% CI [0.57, 0.85], Cohen's d = 1.68, $t(37) = 10.38$, $p < 0.001$) in ADV (Increased SP vs. Decrease SP: 95% CI [0.18, 0.49], $t(37) = 4.44$, Cohen's d = 0.72, $p < 0.001$). And, self-payoff decreases evoked stronger neural responses (0.76 ± 0.09 μV, 95% CI [0.58, 0.94], Cohen's d = 1.42, $t(37) = 8.73$, $p < 0.001$) than self-payoff increases (0.64 ± 0.07 μV, 95% CI [0.50, 0.78], Cohen's d = 1.44, $t(37) = 8.91$, $p < 0.001$) in DIS (Decreased SP vs. Increase SP: 95% CI [0.02, 0.23], $t(37) = 2.44$, Cohen's d = 0.40, $p = 0.04$, Figure 4E left panel). For the late WOI, self-payoff decreases (1.12 ± 0.11 μV, 95% CI [0.90, 1.34], Cohen's d = 1.63, $t(37) = 10.07$, $p < 0.001$) evoked stronger neural responses than self-payoff increases (0.79 ± 0.09 μV, 95% CI [0.61, 0.97], Cohen's d = 1.42, $t(37) = 8.76$, $p < 0.001$) in DIS (Decreased SP vs. Increased SP: 95% CI [0.17, 0.47], Cohen's d = 0.69, $t(37) = 4.26$, $p < 0.001$), but there were no such differences of neural responses between self-payoff decreases (0.87 ± 0.10 μV, 95% CI [0.66, 1.07], Cohen's d = 1.36, $t(37) = 8.36$, $p < 0.001$) and increases (0.99 ± 0.12 μV, 95% CI [0.75, 1.23], Cohen's d = 1.31, $t(37) = 8.10$, $p < 0.001$) in ADV (Decreased SP vs. Increased SP: 95% CI [-0.31, 0.06], Cohen's d = - 0.22, $t(37) = - 1.36$, $p$ = 0.36, Figure 4E right panel).

*Visualization for the relationship between individual differences in altruistic preferences and neural processing of other-payoffs*

To visualize the specific origin of the neural effect identified in the individual-difference analysis of other-payoff processing, we tested the sign and time-course of the corresponding ERP components in the two groups. This revealed that less altruistic participants responded with a more negative early ERP component to increases in other-payoff. To perform this analysis, we categorized trials in the DIS context into trials in which the $2^{nd}$ option increased other-payoff (Increased OP) and trials in which the $2^{nd}$ option decreased other-payoff (Decreased OP) and extracted average ERP waveforms for each other-payoff change level and group. We found that in the less altruistic group, ERP waveforms were negatively correlated with the increase of other-payoff (Increased OP: -0.39 ± 0.20 μV, 95% CI [-0.81, 0.02], Cohen's d = -0.46, $t(18) = -2.00$, $p = 0.06$), and this correlation was significantly stronger than the corresponding correlation (Decreased OP: -0.07 ± 0.18 μV, 95% CI [-0.44, 0.31], Cohen's d = -0.08, $t(18) = -0.36$, $p = 0.72$) with the decrease of other-payoff (Increased OP vs Decreased OP: 95% CI [-0.54, -0.12], Cohen's d =

-0.77, $t(18) = -3.36$, $p = 0.004$). For the more altruistic group, no such effects were observed (Increased OP: -0.06 ± 0.17 μV, 95% CI [-0.41, 0.30], Cohen's d = -0.08, $t(18) = -0.33$, $p = 0.75$; Decreased OP: -0.14 ± 0.16 μV , 95% CI [-0.48, 0.20], Cohen's d = -0.20, $t(18) = -0.89$, $p = 0.39$; Increased OP vs Decreased OP: 95% CI [-0.06, 0.23], Cohen's d = 0.29, $t(18) = 1.25$, $p = 0.23$, Fig. S11).

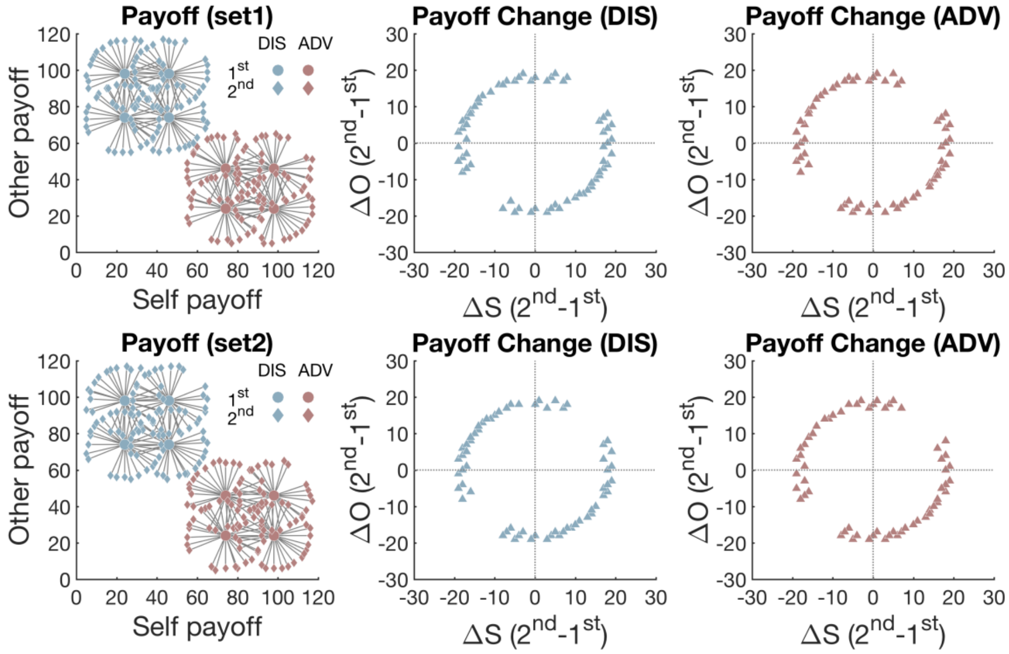


**Fig. S1. Payoff schedule.**

There were two inequality contexts in the task: disadvantageous inequality (DIS) and advantageous inequality (ADV). In the left panel, each dot represents one allocation option and each gray line represents one pair of options that was presented to participants. Blue dots are options in DIS and pink dots are options in ADV. Dots in the center of the circle are the $1^{st}$ options, and diamond dots are the $2^{nd}$ options. Middle and right panels show the distributions of the self-/other-payoff changes between the $2^{nd}$ and the $1^{st}$ option ($\Delta S$ and $\Delta O$) in DIS and ADV, respectively. These two sets of payoff matrix (top panel and bottom panel) have the same reference options and similar distributions of alternative options. By having such payoff matrices of all trials, we matched self-/other-payoff differences and the resulting absolute levels of inequality across both contexts and also across the $2^{nd}$ and the $1^{st}$ options. This allowed us to compare choices and response times, as well as neural processing of different choice features (self- and other-payoff, inequality) between the two contexts.

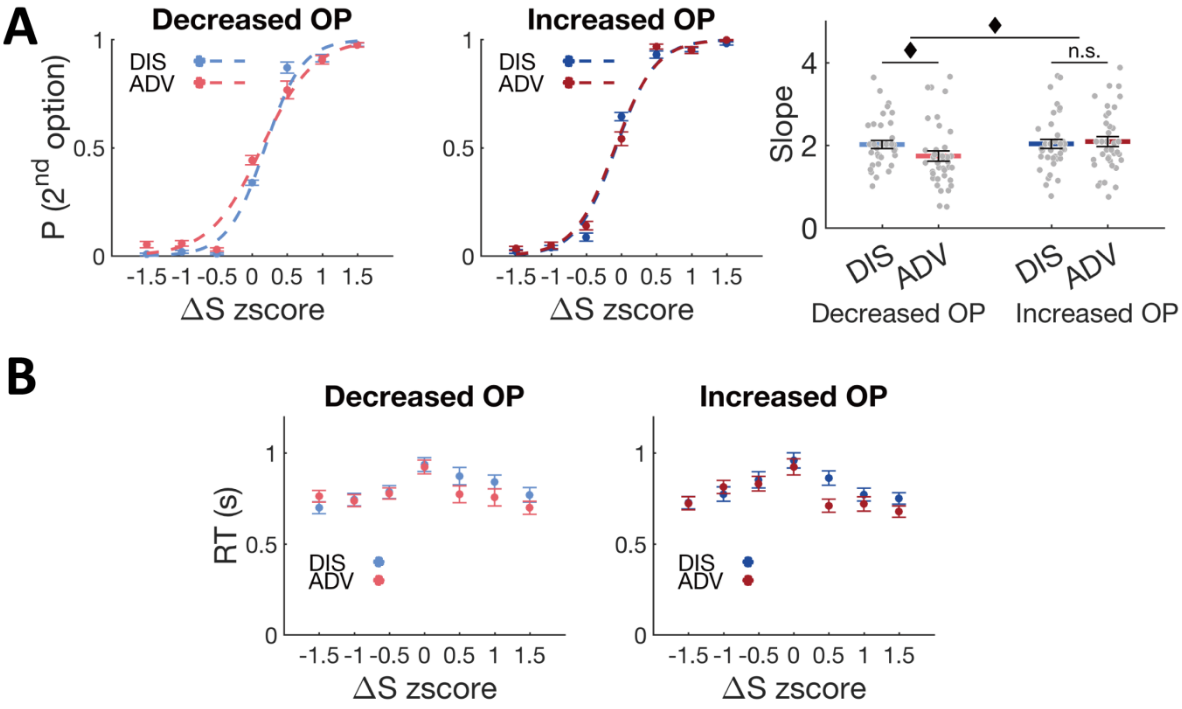


**Fig. S2. Model-free behavioral results.**
(A) Psychometric functions show the effects of ΔS (self-payoff change between the 2nd option and the 1st option) on probability to choose the 2nd option across contexts (CON) and other-payoff change levels (left and middle panels); right panel shows the interactive effect between context and other-payoff change over the slope of the psychometric functions in the middle two panels. Decreased OP, trials in which the 2nd option decrease other-payoff; Increased OP, trials in which the 2nd option increase other-payoff. (B) Plots show the effects of ΔS on RTs across Decreased (Left panel) and Increased OP trials (right panel). OP, other-payoff. ♦, $p < 0.1$.

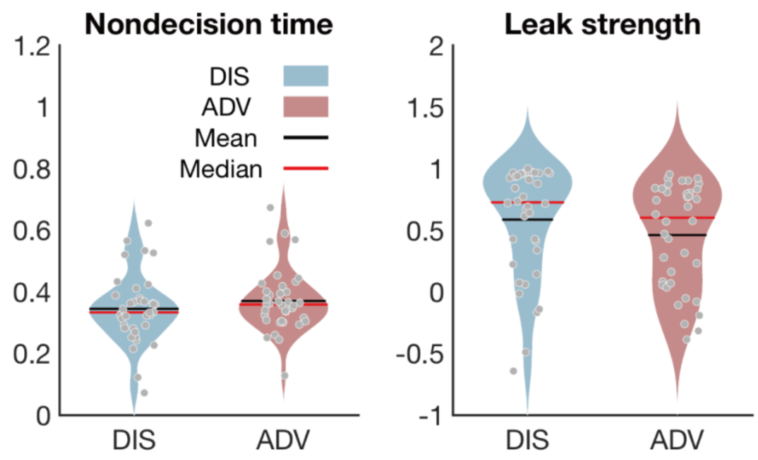


**Fig. S3. OU model parameters in the two inequality contexts.**
Nondecision time and leak strength did not differ between DIS and ADV contexts. Each grey dot represents one participant, and the violin plots represent the distribution. In line with our expectations, nondecision time and leak strength, as two parameters of no-interest, do not contribute to the difference in decision processes of altruism between different inequality contexts.

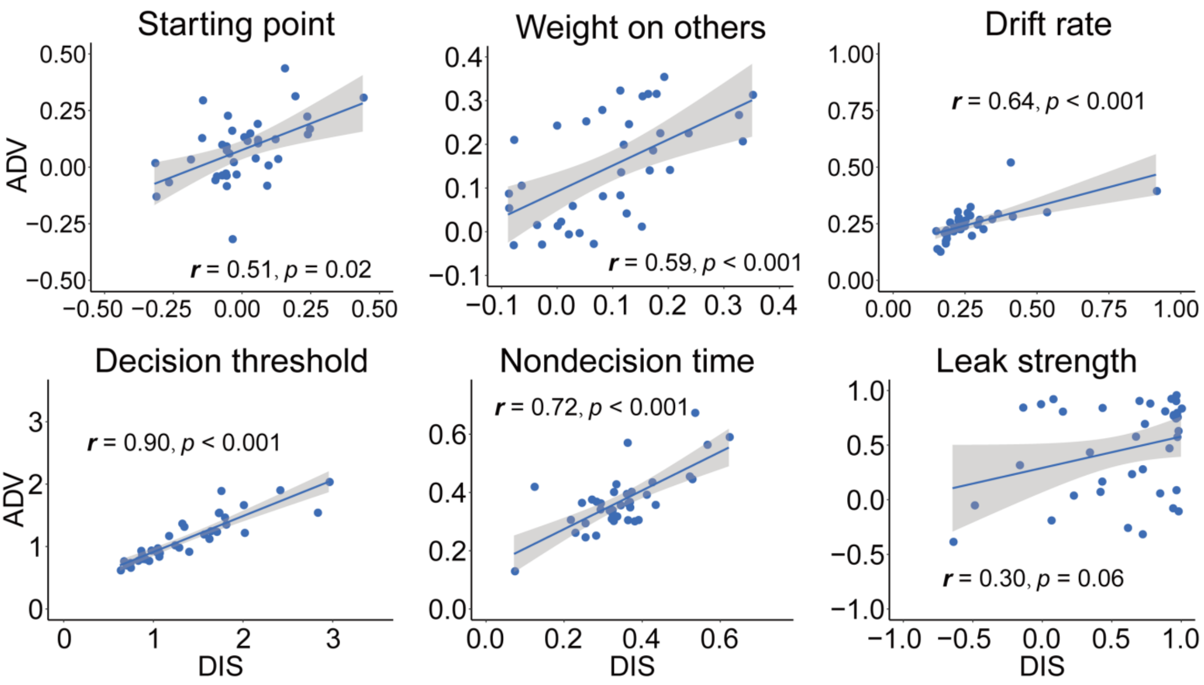


**Fig. S4. Correlations of OU parameters between ADV and DIS contexts.** All fitted parameters are correlated across the two equality contexts, suggesting use of a comparable decision mechanism.

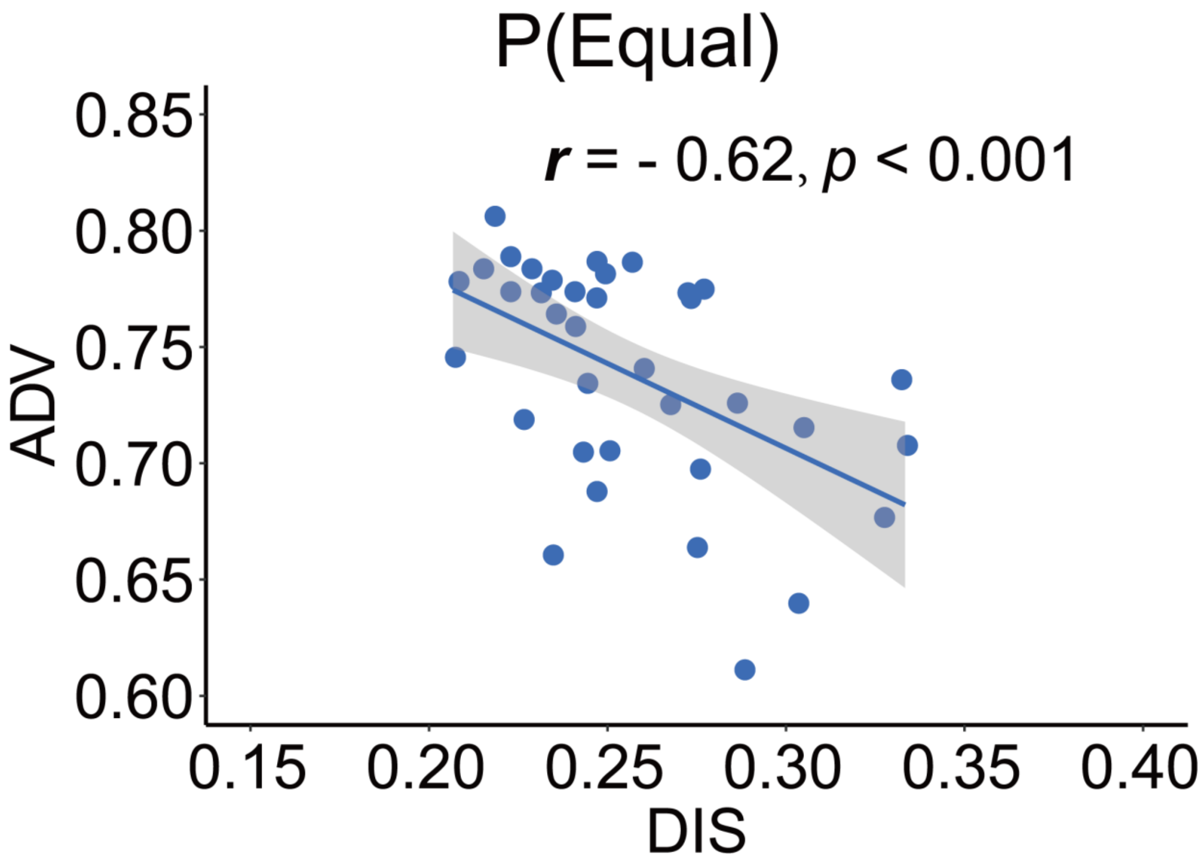


**Fig. S5. Correlation of the probabilities to choose the more equal option in the ADV and DIS contexts.** Significant correlation of the probability to choose the more equal option between ADV and DIS also implied employment of a similar choice mechanism across contexts.

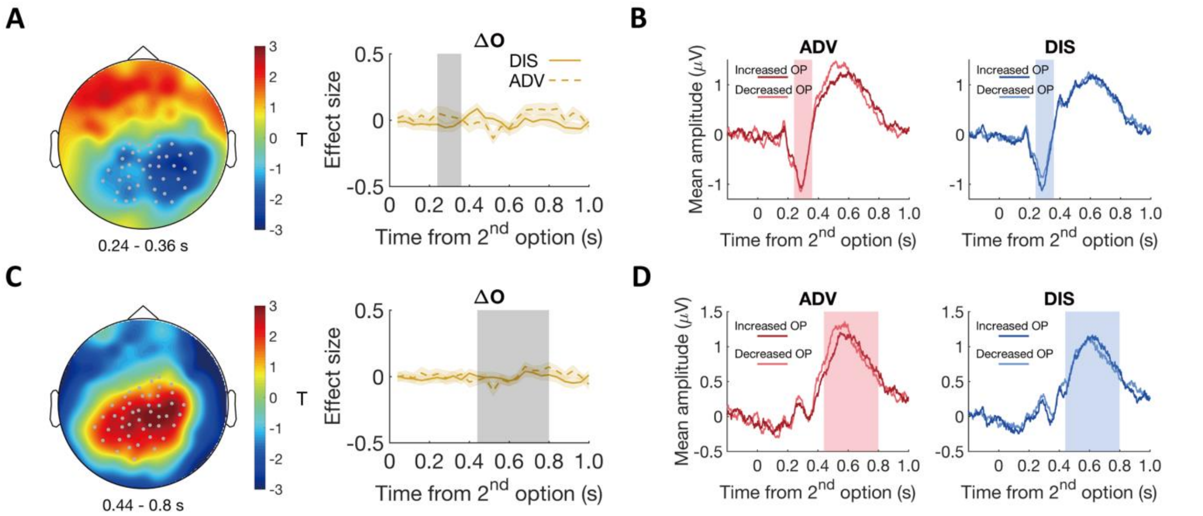


**Fig. S6. Stimulus-locked ERP control analysis of other-interest by context.** To confirm the specificity that context effects in inequality-related neural processing were mainly evident in ERP correlations with self-payoff. We examined the $\beta_2$ (coefficient over other-payoff change ($\Delta O$)) maps, and revealed no ERP components that showed a significant effect of $\Delta O$ in either context, and no differential effects of $\Delta O$ across contexts. Here, we further examine temporal dynamics of the parametric effect strengths of other-payoff change ($\Delta O$) in the specific clusters identified for the context-dependent effects of self-payoff change ($\Delta S$) in the analyses reported in the main text, and revealed no significant effects. (A) & (C) Left panels, topographic scalp distributions showing the significant interaction effects of self-payoff ($\Delta S$) by context in the time windows of ~240 – 360 ms (A) and ~440 – 800 ms (C). Grey dots highlight channels which survive the statistical threshold. (A) & (C) Right panels, temporal dynamics of the parametric effect strengths of other-payoff ($\Delta O$) in the identified clusters related to self-payoff ($\Delta S$) in left panels. Grey shaded areas indicate the durations of the significant effects of self-payoff ($\Delta S$), and colored shaded areas indicate $\pm$ 1 SEM in (A) and (C). (B) & (D) Average ERP waveforms for the effect of other-payoff (ΔO) by context derived from clusters shown in (A) & (C) left panels. In both the time window of ~ 240 to 360 ms after stimulus onset (B, marked by pink and blue shaded areas) and the time window of ~ 440 to 800 ms after stimulus onset (D, marked by pink and blue shaded areas), no significant effect of other-payoff (ΔO) is observed in the ERP waveforms. Pink and blue shaded areas indicate the durations of the significant effects of self-payoff change ($\Delta S$) by context in (B) & (D). Increased OP, trials in which the 2nd option increases other-payoff; Decreased OP, trials in which the 2nd option decreases other-payoff.

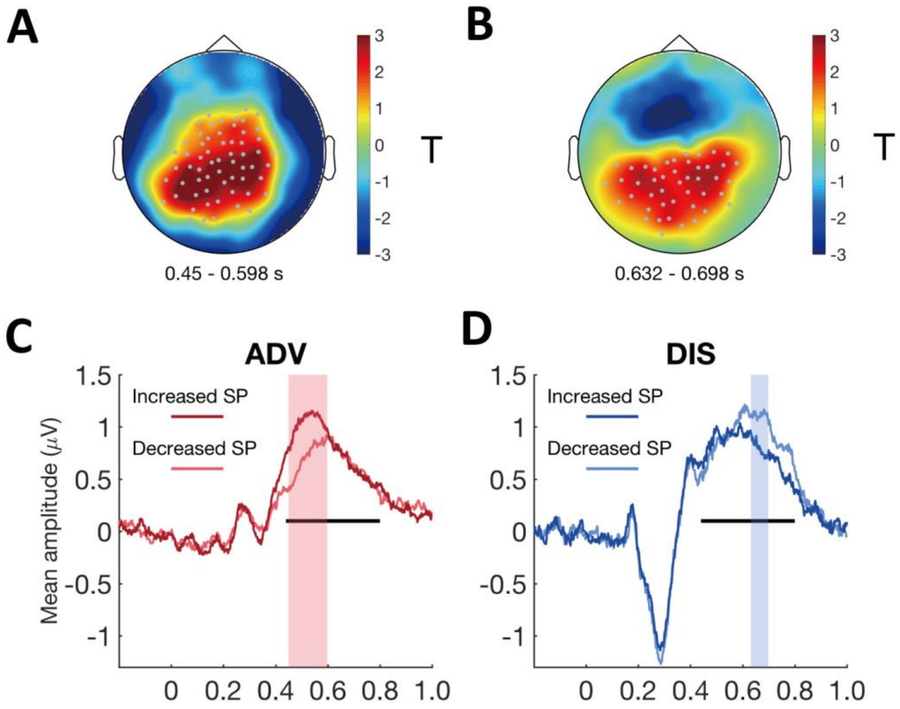


**Fig. S7. Whole brain analyses suggest that the ΔS effect on ERP responses occurs earlier in the ADV than in the DIS context.** The spatially and temporally unbiased whole-brain analysis of the ERP data (not constraining signals to the cluster identified in the initial regression analysis) revealed the same temporal differences between contexts as shown in the main text. These timing differences suggest that the different behavior in the two inequality contexts may relate to a temporally distinct pattern of how neural processing is focused on the inequality level of the choice options, which mainly depends on individuals' assessments of differences in self-payoffs. (A & B) Topographic scalp distributions of the interaction effect between self-payoff change (ΔS) and context at two temporally separate time window. Grey dots highlight channels which survive the threshold. (C) Average ERP waveforms for the effect of ΔS in ADV during the windows of ~450 – 598 ms. (D) Average ERP waveforms for the effect of ΔS in DIS during the windows of ~632 – 698 ms. Black lines indicate the duration of significant interaction between context and self-payoff change, and colored shaded areas indicate the durations of the significant interaction effect of self-payoff change and context in (B) & (D). Increase SP, trials in which the 2$^{nd}$ option increase self-payoff relative to the 1$^{st}$ option; Decrease SP, trials in which the 2$^{nd}$ option decrease self-payoff relative to the 1$^{st}$ option.

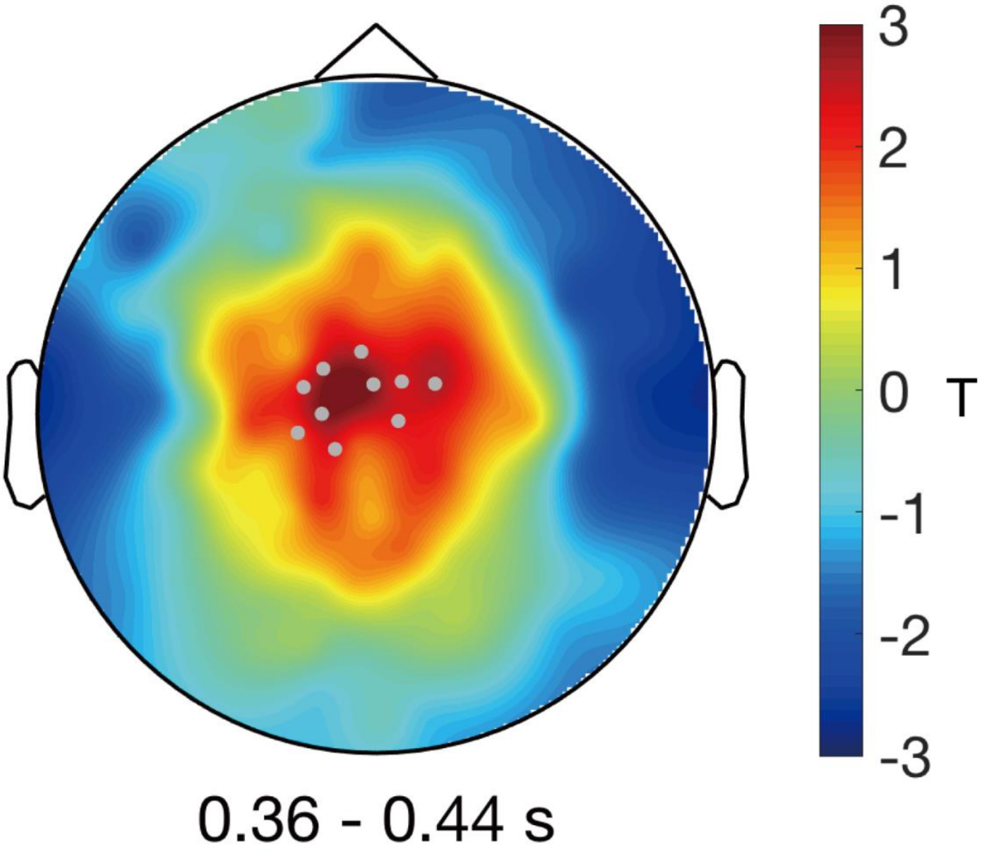


**Fig. S8. Correlation analysis between the model-parameter weight on others' payoffs and neural responses to other-payoff change (ΔO) in the DIS context.** In the main text, we show that for a centrofrontal cluster and the time window of ~320 to 400 ms after stimulus onset, larger increases of other-payoff were associated with a more negative ERP response in the less versus more altruistic group (median-split the weight on others' payoffs (ω),Figure 5A & 5B). A continuous correlation analysis over all participants reveals a similar temporal-spatial cluster as that displayed in Figure 5A, suggesting that the more altruistic a participant, the less negative the responses (in the time window of ~360 to 440 ms after stimulus onset) to larger increases of other-payoff.

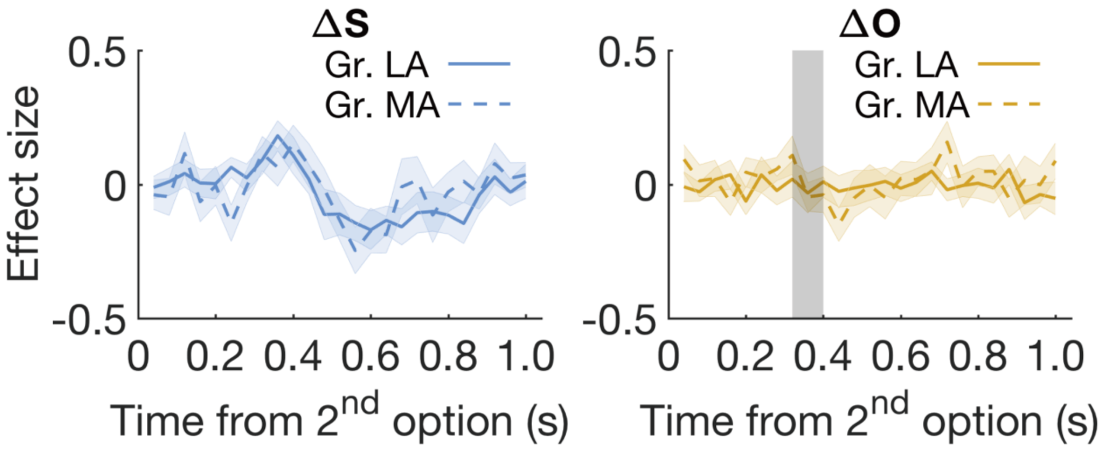


**Fig. S9. Stimulus-locked ERP analysis of other-interest by altruistic preferences.** Participants were divided into two groups (MA vs LA) based on median-split of the starting point $\beta$ parameter in the DIS context (similar to the corresponding analyses based on ω reported in the main text). Temporal dynamics of the parametric effect strengths of self-payoff ($\Delta S$, left) and other-payoff ($\Delta O$, right) in the cluster identified in the individual difference analysis of neural processing of other-payoff ($\Delta O$) based on weight on others. Colored shaded areas indicate $\pm$ 1 SEM. Grey shaded area indicates the duration of the significant effect identified in the individual difference analysis of the OU parameter of weight on others. Gr., group. These findings indicated that the individual difference effect of neural processing of ΔO was specific for ω (weight placed on others' payoffs), and was not evident for $\beta$ (starting point).

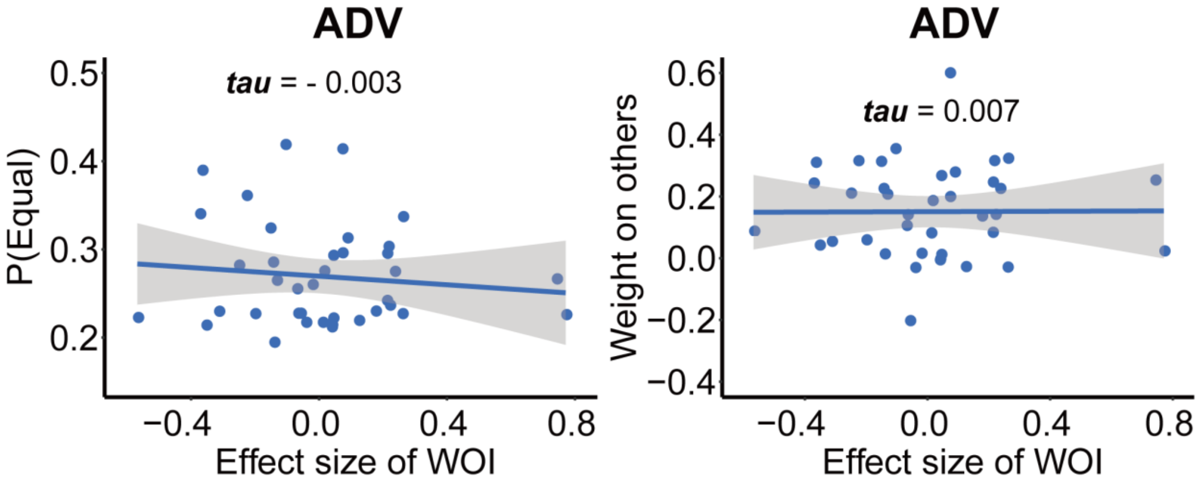


**Fig. S10. Correlations between the effect strengths of other-payoff (ΔO) in ADV (from the cluster identified in the individual difference analysis in DIS) and the probability to choose the more equal option (left), and the OU parameter weight on others (right) in the ADV context.** All these correlations are insignificant, showing the specificity of the effects for the DIS context.

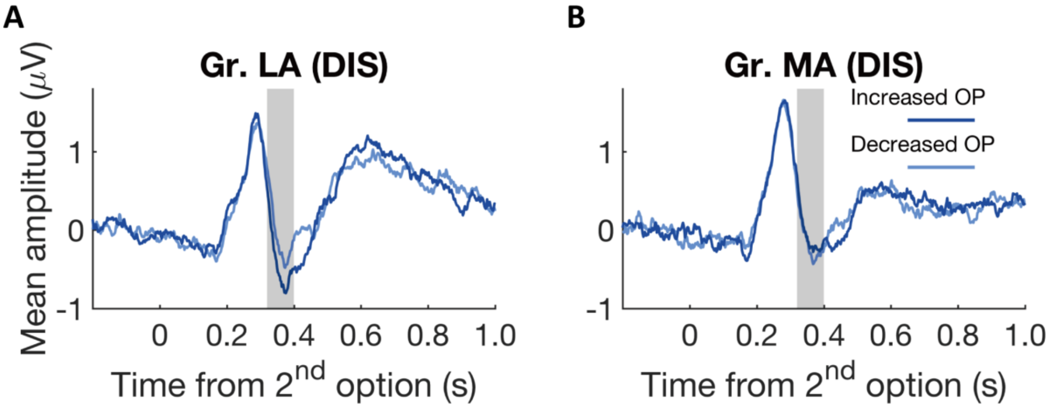


**Fig. S11. Individual differences relate to differential neural processing of other-payoffs in the stimulus-locked ERP analysis**. Average ERP waveforms for the parametric effect of other-payoff change ($\Delta O$) in each group during the windows of ~320 – 400 ms. Increased other-payoff (OP) was associated with a stronger negative-going response than decreased other-payoff (OP) for the less altruistic group (A), but not for the more altruistic group (B). Increased OP, trials in which the 2$^{nd}$ option increase other-payoff; Decreased OP, trials in which the 2$^{nd}$ option decrease other-payoff. Gr., group; MA, more altruistic group; LA, less altruistic group.

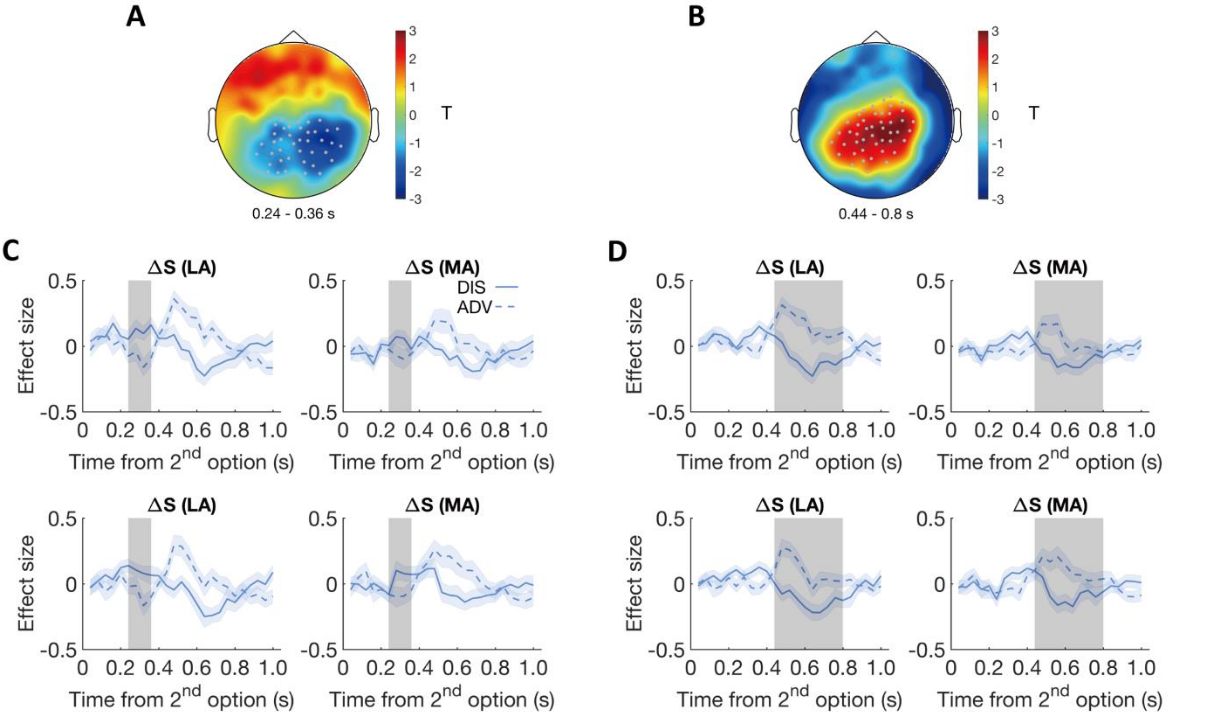


**Fig. S12. Parametric effects of ΔS in the more altruistic (MA) and the less altruistic (LA) groups.** To confirm that individual differences in neural processing of choice-relevant information are unrelated to the context effects, we also compared the more- versus less-altruistic groups in terms of how they processed self-payoff. This showed that irrespective of the grouping parameter (i.e., weight on others or starting point), more altruistic and less altruistic groups exhibited similar neural processing of self-payoff differences in those identified clusters. (A) & (B) Topographic scalp distributions of significant Δ*S* effects by context in the time windows of ~240 – 360 ms (A) and ~440 – 800 ms (B) as shown in Figure 4A and 4C. (C) For the early effect (~240 – 360 ms), MA and LA groups categorized based on weight on others (upper panels) and based on starting point (lower panels) both showed similar dynamic effects as shown in Figure 4A. (D) For the late effect (~440 – 800 ms), MA and LA groups categorized based on weight on others (upper panels) and based on starting point (lower panels) also showed similar effects as shown in Figure 4C. Grey dots highlight channels which survived the threshold in (A) and (B). Blue shaded areas indicate ± 1 SEMs, and grey shaded areas indicate the durations of the significant effects in (C) and (D).

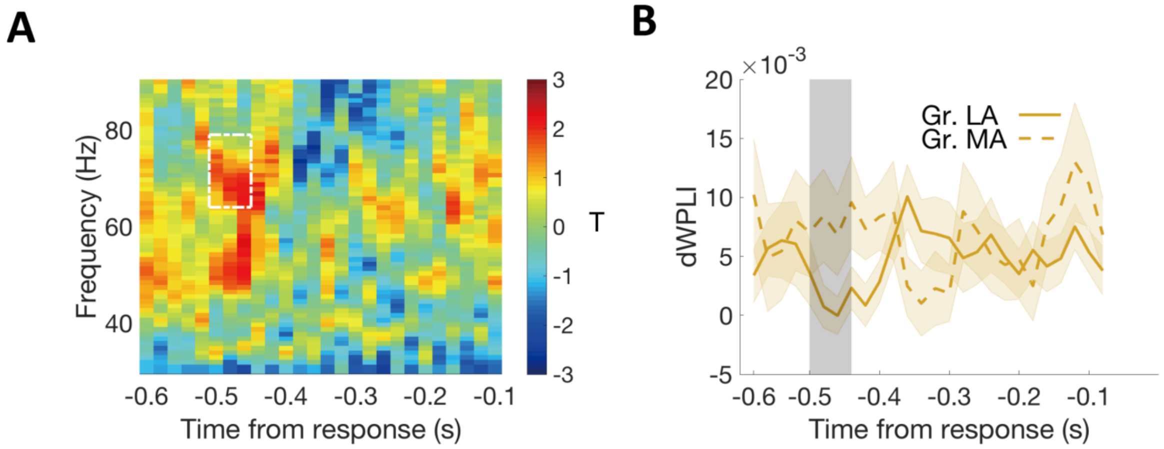


**Fig. S13. Relationship between altruism and frontal parietal synchronization in ADV context.** We do not have a specific hypothesis regarding the relationship between inter-regional synchronization and altruistic preferences for ADV context. Nevertheless, we still tested whether synchronization between centro-frontal regions associated with others' payoff processing and the parietal evidence accumulation regions is associated with altruistic preferences in ADV context. No significant temporal-frequency cluster was identified by this analysis. (A) Heatmap showing T-statistics for the differences between the more altruistic (MA) minus less altruistic (LA) group in phase coupling (dWPLI) between the frontal cluster shown in Figure 5A and the parietal cluster shown in Figure 2D, in the ADV context. White dashed box indicates the temporal-frequency cluster showing significant effect in DIS context. (B) Temporal dynamics of the average dWPLI strengths in the ~64 – 79 Hz frequency range in ADV context. Grey shaded area indicates the duration of the significant effect in DIS context. Colored shaded areas indicate ± 1 SEM. GR, group.

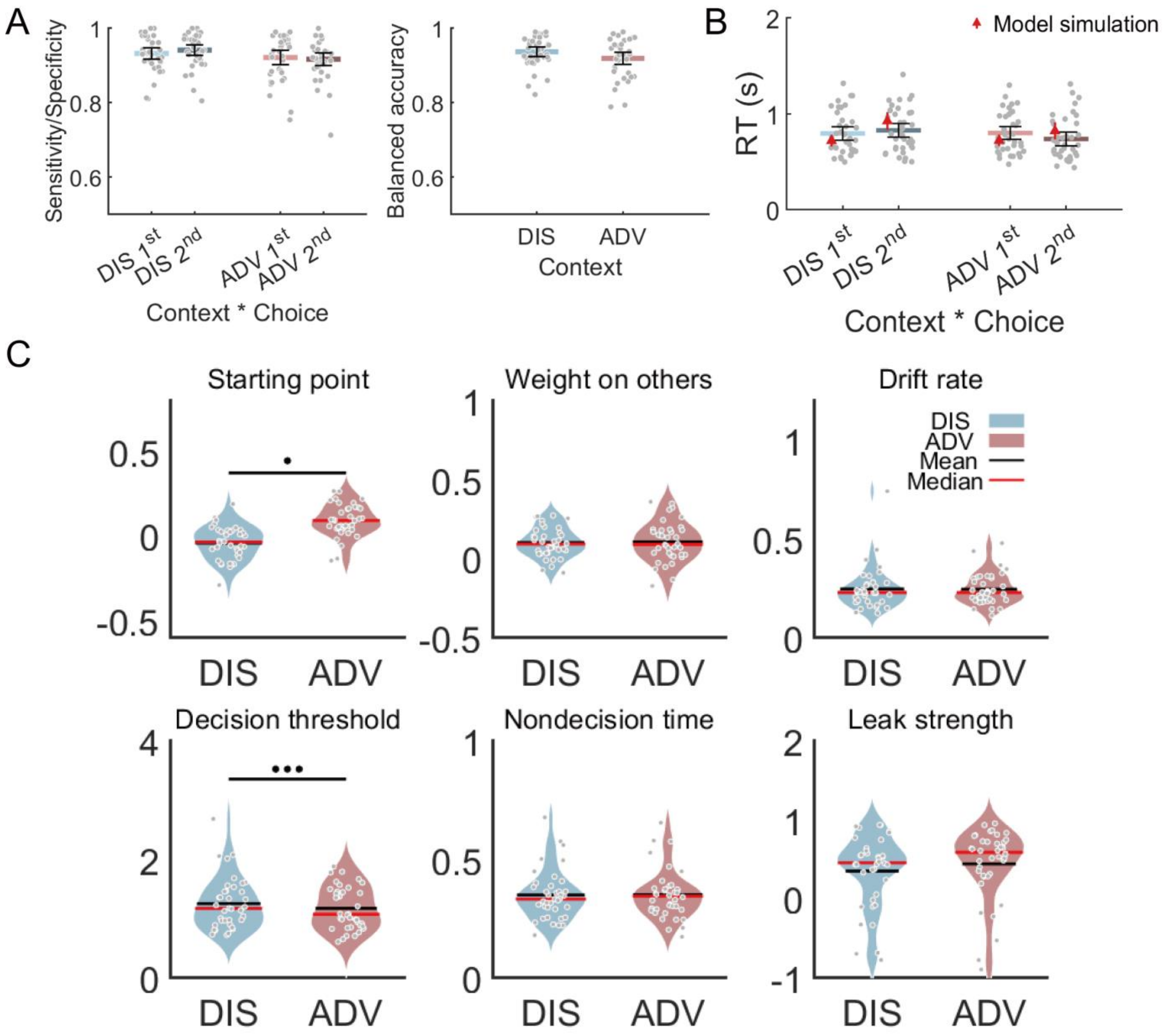


**Fig. S14. Model fits and parameters across inequality contexts**. (A) The OU order model predicts choices across contexts. Left panel: Model performance on Sensitivity of $1^{st}$ \choice /Specificity of $2^{nd}$ choice in each context. Right panel: Model performance on balanced accuracy in each context. DIS, disadvantageous context; ADV, advantageous context. (B) The OU order model recovers RT effects over context and choice in participants' behavioral data. Black error bars display means ± 95% confidence intervals (CIs). Each grey dot indicates one participant. The red triangle dots and error bars represent model simulation mean and 95% CIs. •••, $p < 0.001$. (C) Relative to DIS, ADV drove the starting point towards the $2^{nd}$ offer and reduced decision threshold. Each grey dot represents one participant. More positive starting point is closer to the $2^{nd}$ option.

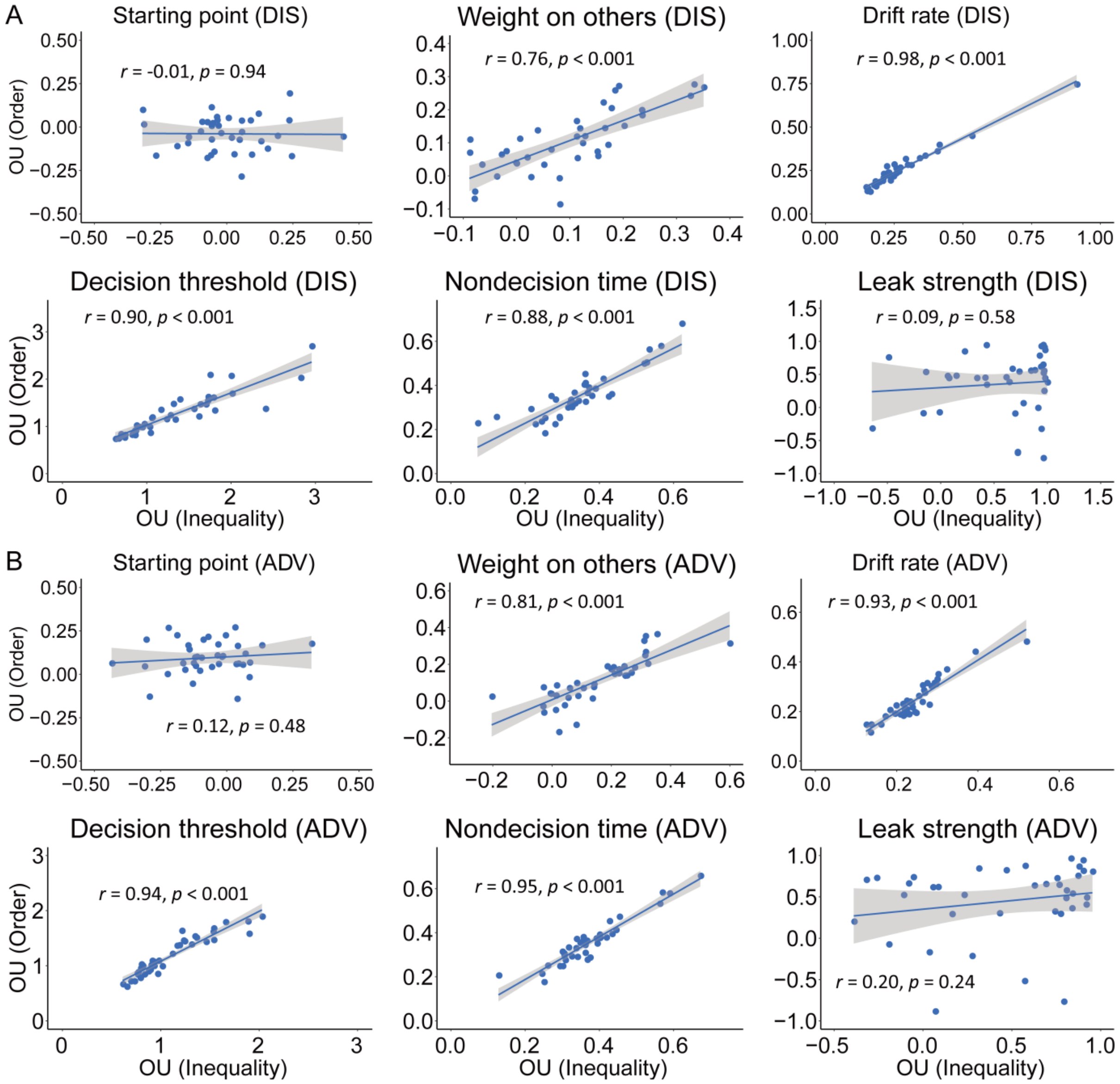


**Fig. S15. Correlations of parameters between the OU order model and the winning OU model (Inequality) in the DIS (A) and ADV (B) contexts.** Weight on others, drift rate, decision threshold, and non-decision time are highly correlated between the two models in both context.

**Table S1. Generalized linear mixed-effects model results of choice data.**

| | Model 1 | | Model 2 | | Model 3 | |
|---|---|---|---|---|---|---|
| Fixed effects | Estimate (95% CI) | p-value | Estimate (95% CI) | p-value | Estimate (95% CI) | p-value |
| Intercept | -0.03 (-0.13 – 0.07) | 0.57 | -0.06 (-0.16 – 0.04) | 0.22 | -0.02 (-0.05 – 0.01) | 0.20 |
| $\Delta S$ | 3.77 (3.65 – 3.89) | < 0.001 | 3.70 (3.59 – 3.82) | < 0.001 | - | - |
| $\Delta O$ | 0.56 (0.51 – 0.61) | < 0.001 | 0.55 (0.51 – 0.60) | < 0.001 | - | - |
| CON | 0.12 (0.06 – 0.19) | < 0.001 | 0.05 (-0.004 – 0.11) | 0.07 | - | - |
| $\Delta S$ * $\Delta O$ | 0.13 (0.01 – 0.24) | 0.03 | - | - | - | - |
| $\Delta S$ *CON | -0.35 (-0.47 – -0.23) | < 0.001 | - | - | - | - |
| $\Delta O$ *CON | -0.02 (-0.07 – 0.03) | 0.50 | - | - | - | - |
| $\Delta S$ * $\Delta O$ *CON | 0.23 (0.11 – 0.35) | < 0.001 | - | - | - | - |
| df | 15,447 | | | | | |
| LL | -3796 | | -3828 | | -10737 | |
| BIC | 7679 | | 7704 | | 21494 | |

$\Delta S$, self-payoff change between the 2nd and 1st option; $\Delta O$, other-payoff change between the 2nd and 1st option; CON, context; CI, confidence interval; df, degree of freedom; LL, log-likelihood; BIC, Bayesian Information Criterion

**Table S2. Generalized linear mixed-effects model results of choice data show that the presentation order (i.e., 1st or 2nd) of the more equal or unequal option does not bias individuals' choices.**

| Fixed effects | Estimate | 95% CI | z-value | p-value |
|---|---|---|---|---|
| Intercept | - 1.07 | -1.35 – -0.79 | -7.41 | < 0.001 |
| $\Delta O$ | 4.83 | 4.61 – 5.05 | 43.65 | < 0.001 |
| CON | -6.32 | -6.60 – -6.04 | -44.58 | < 0.001 |
| Ind_ref | 0.06 | -0.03 – 0.15 | 1.30 | 0.193 |
| $\Delta O$*CON | 0.79 | 0.57– 1.01 | 7.14 | < 0.001 |
| df | 15,486 | | | |
| LL | -5813 | | | |
| BIC | 11684 | | | |

$\Delta S$, self-payoff change between the 2nd and 1st option; $\Delta O$, other-payoff change between the 2nd and 1st option; CON, context; Ind_ref, indicator of the presentation order for the more equal option; CI, confidence interval;df, degree of freedom; LL, log-likelihood; BIC, Bayesian Information Criterion

**Table S3. Linear mixed-effects model results of RT data.**

| Fixed effects | Estimate | 95% CI | t-value | p-value |
|---|---|---|---|---|
| Intercept | 0.79 | 0.72 – 0.86 | 21.98 | < 0.001 |
| $\lvert \Delta S \rvert$ | -0.09 | -0.11 – -0.07 | -7.44 | < 0.001 |
| $\lvert \Delta O \rvert$ | -0.03 | -0.05 – -0.003 | -2.19 | 0.03 |
| CON | -0.03 | -0.04 – -0.02 | -4.28 | < 0.001 |
| $\lvert \Delta S \rvert * \lvert \Delta O \rvert$ | -0.004 | -0.02 – 0.01 | -0.49 | 0.62 |
| $\lvert \Delta S \rvert$*CON | 0.04 | 0.02 – 0.06 | 3.31 | < 0.001 |
| $\lvert \Delta O \rvert$*CON | 0.03 | 0.01 – 0.06 | 2.81 | 0.005 |
| $\lvert \Delta S \rvert * \lvert \Delta O \rvert$*CON | -0.01 | -0.02 – 0.004 | -1.45 | 0.15 |
| df | 15,447 | | | |
| LL | -4184 | | | |
| BIC | 8464 | | | |

$\Delta S$, self-payoff change between the 2nd and 1st option; $\Delta O$, other-payoff change between the 2nd and 1st option; CON, context; CI, confidence interval;df, degree of freedom; LL, log-likelihood; BIC, Bayesian Information Criterion

**Table S4. Bounds of OU parameters.**

| Parameters | Lower bound | Upper bound |
|---|---|---|
| $\alpha$ (Decision threshold) | 0.6 | 3 |
| $\beta$ (Starting point) | -2 | 2 |
| $\kappa$ (Drift rate) | -1 | 1 |
| $\omega$ (Weight on others) | -1 | 1 |
| $\lambda$ (leak strength) | -2 | 2 |
| τ (non-decision time) | 0.01 | 1 |

**Table S5. Model comparison results.**

| Model | Parameters | BIC (Mean ± SE) |
|---|---|---|
| OU model (full) | $\alpha_{(c,s)}$, $\beta_{(c,s)}$, $\kappa_{(c,s)}$, $\omega_{(c,s)}$, $\lambda_{(c,s)}$, $\tau_{(c,s)}$ | 4058 ± 46 |
| DDM | $\alpha_{(c,s)}$, $\beta_{(c,s)}$, $\kappa_{(c,s)}$, $\omega_{(c,s)}$, $\tau_{(c,s)}$ | 4136 ± 48 |
| OU model (fixed $\omega$) | $\alpha_{(c,s)}$, $\beta_{(c,s)}$, $\kappa_{(c,s)}$, $\omega_{(s)}$, $\lambda_{(c,s)}$, $\tau_{(c,s)}$ | 4127 ± 45 |
| OU model (fixed $\alpha$) | $\alpha_{(s)}$, $\beta_{(c,s)}$, $\kappa_{(c,s)}$, $\omega_{(c,s)}$, $\lambda_{(c,s)}$, $\tau_{(c,s)}$ | 4133 ± 46 |
| OU model (fixed $\omega$ & $\alpha$) | $\alpha_{(s)}$, $\beta_{(c,s)}$, $\kappa_{(c,s)}$, $\omega_{(s)}$, $\lambda_{(c,s)}$, $\tau_{(c,s)}$ | 4117 ± 46 |

$\alpha_{(c,s)}$, decision threshold; $\beta_{(c,s)}$, starting point; $\kappa_{(c,s)}$, drift rate modulator; $\omega_{(c,s)}$, relative weight on others' payoffs; $\lambda_{(c,s)}$, leak strength; $\tau_{(c,s)}$, non-decision time (nDT); c for conditions (c = DIS for disadvantageous inequality context, c = ADV for advantageous inequality context), s for participants (s = 1, ..., $N_{participants}$). In the last three models, $\alpha_{(s)}$ and $\omega_{(s)}$ assumed the same decision threshold and/or weight on others' payoff across contexts for each participant. SE, standard error.
